# Obtaining Increased Functional Proteomics Insights from Thermal Proteome Profiling through Optimized Melt Shift Calculation and Statistical Analysis

**DOI:** 10.1101/2022.12.30.522131

**Authors:** Neil A. McCracken, Hao Liu, Avery M. Runnebohm, H.R. Sagara Wijeratne, Aruna B. Wijeratne, Kirk A. Staschke, Amber L. Mosley

## Abstract

Thermal Proteome Profiling (TPP) is an invaluable tool for functional proteomics studies that has been shown to discover changes associated with protein-ligand, protein-protein, and protein-RNA interaction dynamics along with changes in protein stability resulting from cellular signaling. The increasing number of reports employing this assay has not been met concomitantly with advancements and improvements in the quality and sensitivity of the corresponding data analysis. The gap between data acquisition and data analysis tools is even more apparent as TPP findings have reported more subtle melt shift changes related to protein post-translational modifications. In this study, we have improved the Inflect data analysis pipeline (now referred to as InflectSSP, available at https://CRAN.R-project.org/package=InflectSSP) to increase the sensitivity of detection for both large and subtle changes in the proteome as measured by TPP. Specifically, InflectSSP now has integrated statistical and bioinformatic functions to improve objective functional proteomics findings from the quantitative results obtained from TPP studies through increasing both the sensitivity and specificity of the data analysis pipeline. To benchmark InflectSSP, we have reanalyzed two publicly available datasets to demonstrate the performance of this publicly available R based program for TPP data analysis. Additionally, we report new findings following temporal treatment of human cells with the small molecule Thapsigargin which induces the unfolded protein response (UPR). InflectSSP analysis of our UPR study revealed highly reproducible target engagement over time while simultaneously providing new insights into the dynamics of UPR induction.

## INTRODUCTION

The biophysical based Cellular Thermal Shift Assay (CETSA®) and Thermal Proteome Profiling (TPP) assays have been used for nearly a decade to study biochemical phenomena in the cellular context.^1, 2^ Since the initial TPP report,^1^ research groups have leveraged this workflow to identify targets of small molecules and have offered an approach for target and/or off-target identification for drug discovery. Additionally, it has been clearly shown that TPP studies can be used to understand the functional proteome of different cellular states. Recent studies have used TPP to observe: protein stability differences across cell cycle,^3, 4^ protein complex stability,^5^ thermal stability of proteins across evolution,^6^ RNA-protein interactions,^7^ viral infection^8^, and post-translational modifications.^9–12^ Altogether, these studies illustrate the far-reaching potential for TPP to acquire functional proteomics data giving insights into biochemical and biophysical changes that occur in cells and tissues under different experimental conditions. Investigation of protein stability in its native environment has the potential to gain deeper understanding of signaling, protein modifications, and relationships within the cell that have not been observed with prior methods.

Melt curve analysis in TPP is a multi-step process that has been organized in a select group of reported approaches. Algorithms that have been reported by the scientific community are TPP-TR, MSstatsTMT, and Inflect.^13–15^ Advances in TPP data analysis to date, however, have yet to fully address computational challenges in finding reproducible and significant melt shift changes from TPP datasets. One aspect of TPP that has lagged behind is the development of computational tools to determine which melt shifts are “statistically significant”. Variance in quantified protein abundance across temperatures can be compound through the multistep workflow. Changes in stability caused by protein phosphorylation changes, for instance, has been reported by several groups to have limited detectability by TPP.^9–12^ These gaps could be addressed by using additional statistical methods and quality control criteria in the melt calculation pipeline. While some reports have hinted at the utility of statistical methods in the TPP melt shift analysis,^16^ in-depth analysis on how quality control limits affect the output of the data analysis need to be performed. Here we will introduce an updated version of our previously reported program Inflect and have also assessed its utility.^13, 17^ In order to improve the sensitivity and selectivity of the computational workflow for biologically-relevant proteins, we have incorporated a z-score and p-value based assessment into the next version of Inflect, InflectSSP. We have also investigated the relative use of quality control variables in the computational analysis pipeline which include number of peptide spectrum matches or unique peptides in the MS experiment along with the melt curve R^2^. We have also incorporated bioinformatic tools into our InflectSSP program to increase the utility in validating findings from an experiment. While the findings from a TPP discovery experiment may not fully leverage annotated findings, it may be valuable to utilize known protein relationships to reinforce TPP experimental findings. In this work, InflectSSP was used to reanalyze two publicly available data sets from Kalxdorf *et al.* and Sridharan *et al.* to validate its utility and provide benchmarking against other established TPP analysis methods.^18, 19^ Additionally, we designed a temporal experiment through cell treatment with a small molecule inducer of the unfolded protein response (UPR) that we reasoned should cause multiple types of simultaneous changes in protein homeostasis. Overall, our findings show that InflectSSP has increased sensitivity and selectivity for identification of likely changes in the functional proteome while providing multiple statistical metrics including p-value with *post hoc* correction that can be used for cutoff-based analysis of any individual dataset.

## EXPERIMENTAL PROCEDURES

### Data Analysis

TPP experiments were analyzed using InflectSSP as described in the main text. LC-MS/MS data was analyzed using Proteome Discoverer. Bioinformatics analysis was done using annotations from STRING and DAVID databases.^20, 21^ For DAVID analysis a background set of proteins was used that consisted of the list of proteins observed in the respective mass spectrometry experiments.

### Publicly Available Data Sets

Two independent data sets were used in our case study. Data from Sridharan et al.^19^ and Kalxdorf et al.^18^ consist of normalized abundance values at each temperature in the thermal gradients along with search outputs including number of peptide spectrum matches (PSM) and unique peptides (UP). In the case of the dataset from Kalxdorf et al., data was collected in an experiment where K562 cells were treated with 1 mM Dasatinib (protein tyrosine kinase inhibitor) for 60 minutes with the goal of identifying cell surface proteins. Three replicate data sets were used for this analysis. In the case of the Sridharan experiment, the data sets from Jurkat cell crude lysates were treated with 2 mM Na-ATP for 10 minutes. Two replicate experiments were used for this analysis from the Sridharan data set. The data from the source publication including abundance for each protein along with their PSM and UP attributes were used for analysis. In the case of each data set, the qupm columns were used to indicate the number of unique peptides while the qusm column was used for the number of spectral matches. Details regarding the file fields used for data are summarized in Supplemental Table S1.

### Statistical Analysis

*In-silico* experiment design and data analysis were conducted using JMP version 16 (SAS Institute, Cary, North Carolina). The JMP DOE design tool was used to select the combination of PSM, UP, R^2^ and p-value limits for the *in-silico* experiment that would provide a full understanding of “main effects” and “interactions”. The results from the analysis (i.e. number of proteins of interest) were analyzed with respect to each of the inputs using the Fit Model tool in the JMP program. Scaled estimates for each term were calculated by the JMP program and compared to each other in order to understand the relative impact of each term (i.e. PSM) on the output (i.e. number of proteins).

### R Analysis

InflectSSP was coded in R programming language and uses the following functions: readxl, data.table, plotrix, tidyr, ggplot2, xlsx, httr, jsonlite, GGally, network, stats, RColorBrewer, svglite.

### Cell Culture and Treatment

HEK293A cells (Invitrogen, Carlsbad, CA) transduced with a lentivirus encoding an ATF4-firefly luciferase transcriptional reporter gene were used for the Thapsigargin treatment experiments. Cells were cultured at 37°C with CO2 at 5% and water for humidification. Medium consisted of Corning Dulbecco’s Modified Eagle Medium (DMEM, 10-013CV) supplemented with 10% fetal bovine serum (FBS). Cells were grown adherently using 10 cm and 15 cm diameter tissue culture plates. Cultures were passaged every 3-4 days to maintain viability and were also periodically checked for absence of mycoplasma contamination. Treatment experiments consisted of removing growth medium and replacing with fresh media supplemented with Thapsigargin, Tunicamycin or DMSO. Thapsigargin experiments used a 1 mM stock of the drug (Sigma T9033-1MG) dissolved in DMSO. Post treatment for 1 hour, the media was aspirated from the plates and cells were washed with 1X PBS prior to either lysis or removal from the plates. Lysis was used for Western Blot experiments while removal of cells from the plates with rubber scraper was used for TPP and PISA experiments; details are described in respective sections.

### Thermal Proteome Profiling (TPP) Experiments

HEK293A-ATF4-luc reporter cells were treated with 1 µM thapsigargin for 1 – 6 hours, or with vehicle (DMSO) as a control in standard growth media. All cell growths and treatments were performed at least twice to obtain biological replicates. Following treatment, cells were rinsed with cold PBS and harvested by scraping and centrifugation at 300 x g for 5 minutes. Cells were washed again with cold PBS, collected by centrifugation at 300 x g for 5 minutes, and flash frozen at -80 °C. Prior to execution of the TPP workflow, cell pellets were removed from the freezer and were then re- suspended in lysis buffer (40 mM HEPES pH 7.5, 200 mM NaCl, 5 mM Beta glycerophosphate, 0.1 mM sodium orthovanadate, 2 mM TCEP, 0.4% NP40, 1X Roche EDTA free mini complete protease inhibitor). Cells were lysed 1.5 mL Micro Tubes (Diagenode) by sonication using a Bioruptor® sonication system (Diagenode) with cycles of 30 seconds/30 seconds off for 15 mins in a 4°C cold water bath. Total protein concentration of each sample was determined by a Bradford protein assay with lysates diluted to a protein concentration of 4 mg / mL for subsequent temperature treatment. Aliquots of the adjusted supernatants (50 μL) were transferred to PCR tubes after which the samples were heated and cooled using a gradient procedure. The heat treatment consisted of 2 minutes at 25° C, 3 minutes at given temperature per gradient, 2 minutes at 25° C, followed by 4° C. Gradient temperatures for TPP experiments consisted of 25.0, 35.0, 39.3, 50.1, 55.2, 60.7, 74.9 and 90.0° C. Heat treated samples were centrifuged at 20,000 rcf for 30 minutes to pellet insoluble protein while the supernatant was reserved and precipitated in 20% Trichloroacetic acid (TCA).

### Sample Preparation for LC-MS/MS and Mass Spectrometry

Dried pellets were resuspended in 8M Urea in 100 mM Tris pH 8.5. Samples were reduced with TCEP and alkylated with chloroacetamide as previously reported.^22^ Reduced and alkylated samples were digested with LysC/Trypsin (Promega) followed by quenching with formic acid. Quenched samples were desalted with Waters C18 columns and then isobarically labeled with Thermo Scientific TMTPro labels (Lot UI292951) as previously reported.^23^

### Search Parameters and Acceptance Criteria (MS/MS and/or Peptide Mass Fingerprint (PMF) data)

There were two total technical replicates of the Thapsigargin datasets using two different LC-MS instruments. In the case of one technical replicate, Nano-LC-MS/MS analyses were performed on an Exploris 480 mass spectrometer (Thermo Scientific) coupled to an EASY-nLC HPLC system (Thermo Scientific). The peptides were eluted using a mobile phase (MP) gradient as follows with 95% phase A (FA/H2O 0.1/99.9, v/v) to 24% phase B (FA/ACN 0.1/80, v/v) for 150 mins, from 24% phase B to 35% phase B for 25 mins and then holding for 5 min at 400 nL/min to ensure elution of all peptides. Nano-LC mobile phase was introduced into the mass spectrometer using a Nanospray Flex Source (Proxeon Biosystems A/S). The heated capillary temperature was kept at 275 °C and ion spray voltage was kept at 2.5 kV using a FAIMS source with a compensation voltage of -50V. During peptide elution, the mass spectrometer method was operated in positive ion mode for 180 mins, programmed to select the most intense ions from the full MS scan using a top speed method. Exploris MS1 parameters include: Microscans 1; MS1 Resolution 60k; automatic gain control (AGC) target 3E6; and Scan range 375 to 1600 m/z. Exploris data dependent MS/MS parameters include: Microscans 1; Resolution 45k; AGC target 2E5; Maximum IT 87 ms; Isolation window 0.7 m/z; Fixed first mass 110 m/z; and HCD normalized collision energy 35.0. The respective data dependent settings were set with parameters: Apex trigger as “-”; Charge exclusion as “1,7,8, >8“; Multiple Charge. States as “all”; Peptide match as “preferred”; Exclude isotopes as “on”; Dynamic exclusion of 30.0 s; If idle “pick others”. The data were recorded using Thermo Xcalibur software (Thermo Fisher).

A technical replicate analysis of the samples was also performed on a Lumos mass spectrometer (Thermo Scientific) coupled to an EASY-nLC HPLC system (Thermo Scientific). The peptides were eluted using a mobile phase (MP) gradient as follows with 94% phase A (FA/H2O 0.1/99.9, v/v) to 25% phase B (FA/ACN 0.1/80, v/v) for 170 mins, from 25% phase B to 80% phase B for 10 mins at 400 nL/min to ensure elution of all peptides. Nano-LC mobile phase was introduced into the mass spectrometer using a Nanospray Flex Source (Proxeon Biosystems A/S). During peptide elution, the mass spectrometer method was operated in positive ion mode for 170 mins, programmed to select the most intense ions from the full MS scan using a top speed method. Lumos MS1 parameters include: Microscans 1; MS1 Resolution 120k; standard automatic gain control (AGC); and Scan range 400 to 1600 m/z. Lumos data dependent MS/MS parameters include: Microscans 1; Resolution 50k; Normalized AGC Target of 250%; Isolation window 0.7 m/z; Fixed first mass 100 m/z; and HCD normalized collision energy 34.0. The respective data dependent settings were set with parameters: Exclude isotopes as “on”; Dynamic exclusion of 60.0 s. The data were recorded using Thermo Xcalibur software (Thermo Fisher).

The resulting RAW files were subjected to protein FASTA database search using Proteome Discoverer 2.4.0.305 (Thermo Scientific, Waltham, MA). The SEQUEST HT search engine was used to search against a human protein database from the UniProt repository containing 20,350 human proteins (2019) and common contaminant sequences such as proteolytic enzymes (FASTA file used available on MassIVE under MSV000090867 and in ProteomeXchange under PXD038752). Specific search parameters used were trypsin as the full proteolytic enzyme, peptides with a max of two missed cleavages, precursor mass tolerance of 20 ppm, and a fragment mass tolerance of 0.5 Da. Minimum and maximum peptide length were set to 6 and 144 respectively with max number of peptides reported at 10. Spectrum matching parameters in the search were set to True for “Use Neutral Loss” for all ions and weight of b and y ions were set to 1 with all others at 0. Max equal and dynamic modifications per peptide were set to 3 and 4 respectively. Static modifications were TMTPro label on lysine (K) and the N-termini of peptides (+304.207 Da). Percolator false discovery rate (FDR) cutoff filtering was set to a strict setting of 0.01 (1% FDR). Total ion abundance values at the protein level were summed from unique peptides and used for quantification for melt shift calculation at the protein level. The mass spectrometry proteomic data have been deposited to the ProteomeXchange Consortium via the MassIVE partner repository with the data set identifier and doi:<u>10.25345/C5VM4325J</u> and login information available upon request prior to publication.

### Experimental Design and Statistical Rationale

The Sridharan experiment analysis used two publicly available biological replicate data sets while the Kalxdorf experiment analysis used three publicly available biological replicate data sets (total number available from each respective source). The Thapsigargin data sets from our group consisted of two biological replicates and this number of experiments was chosen based on the number of temperatures that could be successfully multiplexed using available TMT labels. Each of the three data set used both a condition (treatment) and control (vehicle) to calculate melt shifts. *In-silico* experiment design and statistical data analysis were conducted using JMP version 16 (SAS Institute, Cary, North Carolina). TPP experiments were analyzed using InflectSSP version 1.4.5 (described herein). The InflectSSP program has an optional False Discovery Rate adjustment which can be used by users to increase the specificity of the data set being analyzed. To account for potential biological and workflow variability in data sets analyzed, a false discovery rate calculation was not used in the analysis described herein. The z-score based p-value calculated by the InflectSSP program was deemed acceptable for the sigmoidal data analysis that was being analyzed. Traditional coefficient of determination (R^2^) is also available for use in the InflectSSP program as it is used in the field for describing the adequacy of sigmoidal fit.

## RESULTS

### Initial assessment of InflectSSP workflow

Inflect was developed as an R package for the analysis of thermal proteome profiling experiments. A goal of our ongoing development of the Inflect workflow is to increase the sensitivity and selectivity for changes in protein thermal stability when analyzing TPP (and similar assay) results. Using the output from a TPP experiment, normalized ion abundance values at different temperature treatments are used as input for the melt curve analysis in Inflect to determine the inflection point of the curve as the melt temperature (Tm) of the protein of interest (POI). For development of InflectSSP, additional parameters were developed to consider their respective impact on the output of the melt shift calculation workflow. The InflectSSP data analysis workflow that was used for our assessment is described pictorially in Figure 1. **Step A** in the workflow imports data from the source directory. The user will specify the number of replicate experiments. The current version of InflectSSP allows for the import of multiple experiments which can consist of an unbalanced number of experiment files for condition and control. The normalization step divides each abundance value by the abundance observed at the lowest temperature so that results from separate experiments can be analyzed together. This data normalization is completed at the “protein level” for each protein and for each experiment factor (vehicle or drug). If for instance there is a vehicle treatment and a drug treatment (2 factors) in the 8- temperature heat treatment experiment and a total of 5,000 proteins have been identified, there will be 80,000 abundance values at the end of this “Normalization” step. **Step B** in the algorithm converts the normalized abundance data from the protein level to proteome level. Specifically, the median abundance is measured at each temperature across all factors. In our example, if there are 80,000 abundance values at the end of step A with 8 heat treatments, this “Quantitation” will yield 8 total values. **Step C**, or “Curve Fit 1” determines the three or four parameter log fit coefficients that best describe the variability observed in Step B. The purpose of steps B and C are to describe how well the heat treatment step compared to ideal melt behavior. If for example, there is a subtle increase or decrease for one temperature in the heat gradient (e.g. in the heating block), the curve would depart from sigmoidal shape. A four-parameter log fit (4PL) is used to describe the curve but changes to a three-parameter log fit (3PL) if there are challenges with curve fit convergence by the program. **Step D** is the “Correction” step that adjusts the normalized abundance value at each temperature for each protein based on how well actual values meet predicted values in Step C (“Curve Fit 1”). While Step C is done at the proteome level, Step D is done at the protein level. In this step, a correction factor is first calculated for each temperature based on how much the actual values departed from predicted values in Step C. The correction factor at each temperature is then used for each protein in the experiment. For example, if the actual values were 1% greater than those predicted at 35°C, the normalized abundance values for each individual protein at 35°C would be decreased by 1% percent to allow for normalized abundance values to fit with ideal melt behavior. If there are 5,000 proteins going into Step B and there are 100 proteins excluded due to low PSM or UP, there would be a total of 4,900 proteins going into Step D and therefore 78,400 normalized abundance values being corrected. “Curve Fit 2” in **Step E** is executed for each individual protein in the experiment at the protein level. In our example, 4,900 proteins across two factors would be used to fit 9,800 curves.

**Figure 1.**
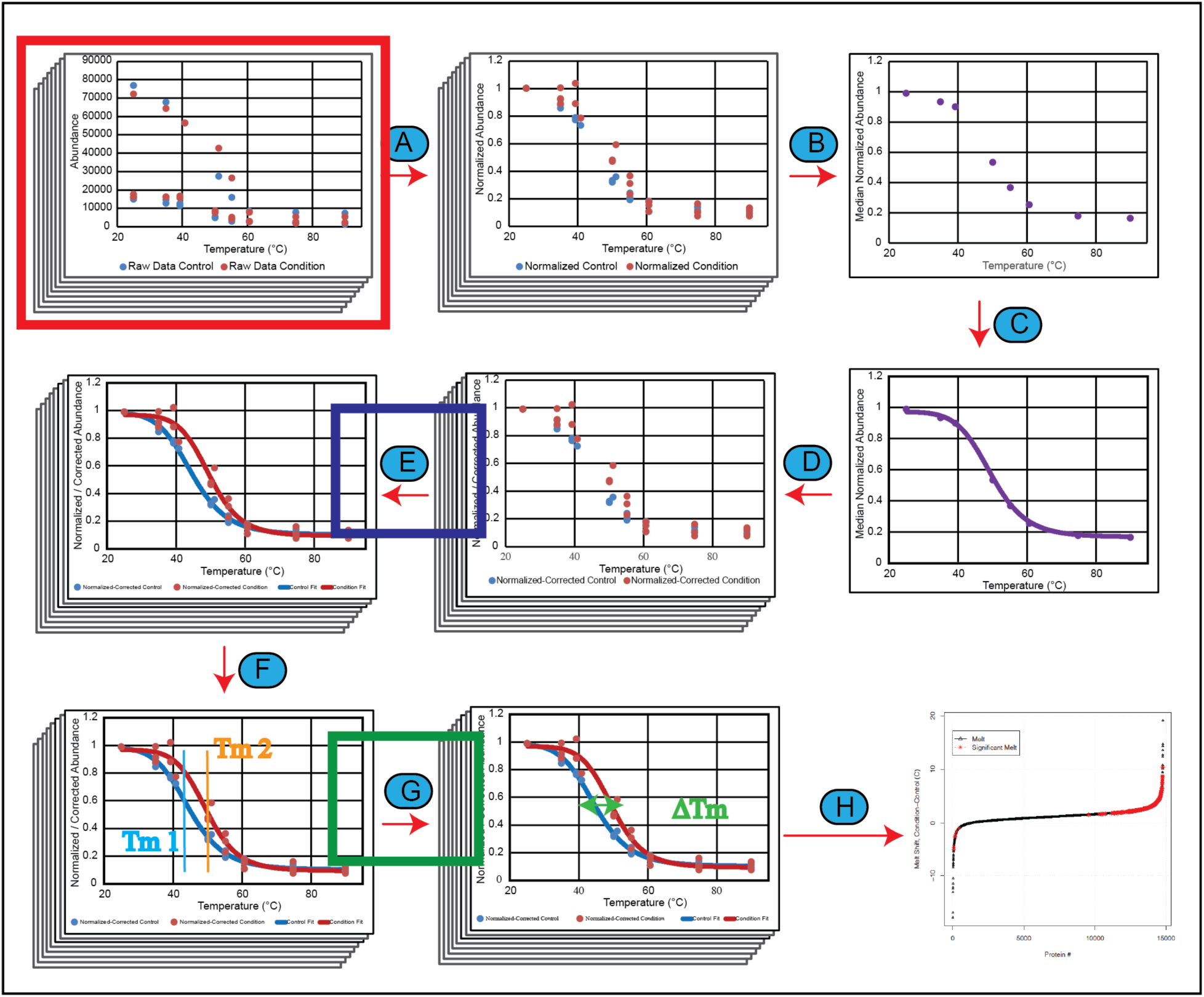
(A) Abundance data from each protein normalized by dividing each abundance value at each temperature by the abundance at the lowest temperature. (B) Median of all abundance values across proteome at each temperature and for each condition and control across all replicates. (C) Curve fitting of proteome data sets. (D) Correction of individual normalized abundance values for each protein based on difference between actual and predicted in previous step. (E) Curve fitting for each protein melt curve based on corrected normalized abundance values. R2 for curve fits are also calculated. (F) Calculation of melt temperature for each condition and control across all proteins. (G) Calculation of melt shift along with melt shift p-value. (H) Result reporting that describes the melt shifts calculated in the experiment with outputs in various formats (plots and tables).Potential locations in the data analysis workflow where quality of the data analysis can be evaluated. The red box indicates the number of peptide spectrum matches (PSM) and number of unique peptides (UP) that are reported with the protein/abundance data from the MS experiment and proteomic search. The blue box represents where the coefficient of determination (R2) is calculated based on the fit of the melt curves. The green box represents where the melt shift p-values are calculated in the workflow with the melt shifts.

Like “Curve Fit 1” in Step C, a 4PL is first used before using a 3PL fitting. If neither set of equations converges in the program, the protein is excluded from further analysis. The melt temperature for each protein is calculated in Step E using the inflection point of the melt curves. The inflection point is defined as the temperature where the 2^nd^ derivative of the fit equation equals 0. In the process of generating the curve fits and the associated inflection point, it is possible that the curve fitting algorithm can converge on a set of optimal parameters. The fit curve, however, may not represent what would be biologically possible. To address this challenge, the 3PL fit is used if a calculated melt temperature is less than or greater than the temperature range used during the heat treatment. An example of how this operation allows for more biologically representative results is shown in Supplemental Figure S1. In this example where a 4PL fit was initially used, the melt is calculated to be 71.3°C but when the 3PL fit was used (to better reflect biological conditions) a more realistic melt of 57.0°C is observed. **Step F** or “Melt Calculation” is completed using the fit curves for each protein. Specifically, the melt temperature for each protein is calculated as the inflection point in the sigmoidal curves that are calculated in the previous step. This process is also linked with the previous “Curve Fit 2.” If the calculated melt temperature is less than the lowest temperature or greater than the highest temperature in the heat treatment, a 3PL will be used. This process has been implemented in this next version of Inflect to avoid artificially large melt shifts that result from melt curve shapes that are not anticipated to reflect biological conditions. The “Melt Calculation” is completed in **Step G** where the control melt temperature is subtracted from the condition melt temperature to determine the magnitude of shifts for each protein in the experiment. In our example to this point, the output of this step would be 4,900 melt shifts. **Step H** consists of generating a rank order or waterfall plot that describes the melt shifts of each rank ordered protein in the experiment.

The workflow used to analyze data from the Kalxdorf et al. and Sridharan et al. data sets is described in the materials and methods section of this report. In the case of the dataset from Kalxdorf et al., data was collected in an experiment where K562 cells were treated with 1 mM Dasatinib (protein tyrosine kinase inhibitor) for 60 minutes with the goal of identifying cell surface proteins. Three replicate data sets were used for this analysis. In the case of the Sridharan experiment, the data sets from Jurkat cell crude lysates were treated with 2 mM Na-ATP for 10 minutes. Two replicate experiments were used for this analysis from the Sridharan data set. The rank order plots of the calculated melt shifts for each protein in these data sets are shown below in Figure 2A and Figure 2B and these panels reflect the wide range of melt shifts in each set. While 10-20% of the protein melt shifts are greater than 2°C and around 4% of the proteins are greater than 5°C (Figure 2C and Figure 2D), it is not easily discernible from melt shifts alone which proteins have a significant melt shift and which are within the variability of the experiment. One possible method for determining significance would be to report proteins with melt shifts that are greater than an absolute limit based on the mean and standard deviation. At the same time, this approach may not allow for selection of proteins with subtle changes. This fact guided our work to provide objective quality control criteria based on statistical metrics to apply cutoffs for selecting proteins of interest in these data sets.

**Figure 2.**
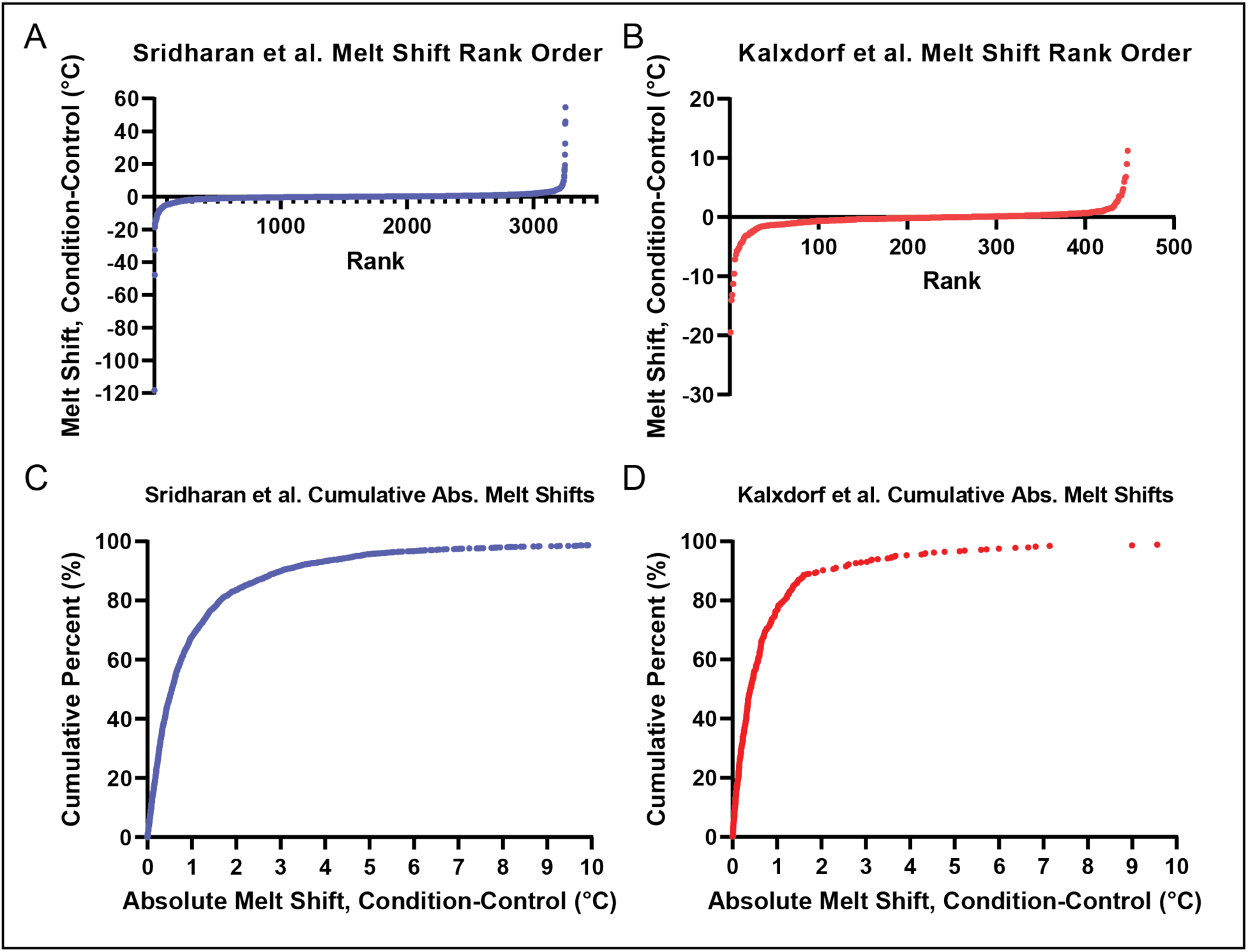
Melt shift rank order plots for (A) Sridharan and (B) Kalxdorf data sets. Pareto chart for absolute melt shift shifts for (C) Sridharan and (D) Kalxdorf data sets. X-axis max for (C) and (D) have been set at 10°C for ease of viewing.

The absolute magnitude of melt shifts in a TPP experiment may not be sufficient for determining which proteins are significantly stabilized or destabilized for functional proteomics interrogation. The reason for this limitation is that the calculated melts may not account for experimental variability. The melt curve inflection point or melt temperature may be distinct, but the variability of the abundance values around the calculated melt from the curve fit may be large enough that the melt temperatures between condition and control are not significantly different from each other. Selection of a protein based on the magnitude of the shift alone would potentially cause the investigator to focus on proteins and pathways that are not actually affected by the experimental conditions. To address this deficiency in the calculation pipeline, we have updated our existing version of Inflect to allow for analysis of biological replicate experiments to determine statistical significance of melt shifts between conditions. Other filters or quality control steps were also added to the analysis pipeline to remove proteins that increase the “noise” of the calculated melt shift. Three steps in the described analysis workflow (Figure 1) were identified as possible opportunities for inserting control criteria. The mass spectrometry and proteomics search (designated by the red box in Figure 1) is one place where criteria can be set. The number of unique peptides (UP) for each protein is one output that is reported by proteomic search algorithms such as Proteome

Discoverer.^a^ This value indicates the number of reported peptide fragments that are unique to a protein in the source proteomics database. Since the sequence of peptides determined from a “bottom-up” mass spectrometry experiment is based on experimental spectra, the number of peptide spectrum matches (PSM) are also reported by the proteomics software. The PSM are the number of spectra from the experiment that match with spectra from the search database within the set cutoff criteria. These two variables offer quality control criteria for screening the performance of the mass spectrometry and associated data search experiment. In our analysis, we used the UP and PSM reported in Step A (Figure 1) to conduct the exclusion of proteins at step D after the proteome curve fit is conducted. These filters allow for exclusion of proteins with low numbers of total identifications independent of their summed ion abundance value. These two filters were inserted prior to the curve fitting steps in Step E since a low UP or PSM value could result in lower confidence in the identity of the protein associated with the peptide(s). The exclusion was set after the overall proteome curve fitting (Step C) to ensure that the abundance values of the peptides still affect the total proteome abundance. The distribution of PSM and UP across the two data sets is shown in Supplemental Figure S2A and Supplemental Figure S2B. Both sets of distributions for both data sets are skewed to low number of total PSMs and unique peptides as is commonly observed for DDA-based bottom-up proteomics experiments. Considering that the multiplexing allowed for by TMT-based isobaric labeling is required for analysis of a high-dimensional TPP dataset with multiple temperature datapoints, this challenge cannot be directly addressed using alternative acquisition strategies such as data independent acquisition which is not compatible with isobaric labeling.

A second step in the analysis workflow where filtering was identified is Step E where the curve fitting is conducted (blue box in Figure 1). One statistical tool that can be used to describe the quality of fit for the melt curve is the coefficient of determination (R^2^). While the coefficient of determination is not necessarily a strong measure of non-linear fit,^24, 25^ it was ascertained that the wide use of this parameter in the scientific community still maintained its utility. This step in the process was identified as a point where variability in the cell culture, heat treatment and MS portions of the experiment could be characterized. This was also a way of incorporating variability from replicate experiments. The distributions of R^2^ for each of the two data sets are shown in Supplemental Figure S2C and reflects the fact that the quality of fit is skewed towards values of 1.

A third point in the overall workflow where possible data filtering was identified is Step G where the melt shifts are calculated (green box in Figure 1). The difference in melt temperature between condition (i.e. treatment) and control (i.e. vehicle) in an experiment is defined as the melt shift. A positive shift is generally interpreted as a protein with increased thermal stability while a negative shift is a decrease in protein thermal stability from experiment conditions. The magnitude and direction of each shift do not provide sufficient information to say whether a shift is greater than experimental variability. Experiment variability can be captured through the execution of biological and technical replicates that are input into the InflectSSP workflow. The melt curves are fit for each protein in Step D (Figure 1) using either four-parameter or three-parameter log fits (4PL or 3PL respectively) depending on the success with analysis convergence. To quantify the signal to noise ratio in these melt shifts, we have established a z-score accompanied by a p-value calculation. The equation in Figure 3A is used by the current version of InflectSSP to calculate a p-value for each protein melt shift. The calculation is based on the difference in melt temperature normalized by the standard error calculated by the nls function. Our z-score uses a 1 standard deviation criteria for the evaluation of the p-value. The definition of this p-value is the likelihood that you would reject the null hypothesis (no difference in melt temperature) when the null hypothesis is true. Supplemental Figure S2D shows melt shift p-value distributions for each of the two data sets and indicates that while the Sridharan data set is skewed toward more statistically significant melts (lower p-value) the Kalxdorf data set has a more even distribution of melt p-values with less statistically significant shifts. Figure 3B shows the distribution of melt shifts that have p-values less than 0.05 and reinforces the fact that melt shifts that meet the p value of < 0.05 are not necessarily large in magnitude, clearly showing the utility of a statistical method that considers the reproducibility of biological replicates. The plots in Figure 3C and Figure 3D show the relationship between the calculated melt shift p-value and the melt shifts across both data sets. These results indicate that while there are a significant number of proteins with high melt shift that have a low p-value, there are also a significant number of proteins that have a low p-value accompanied by a low melt shift. These results suggest that the magnitude of the melt shift is insufficient for determining the significance of the shift as many of the proteins with low p-values also have very small shifts.

**Figure 3.**
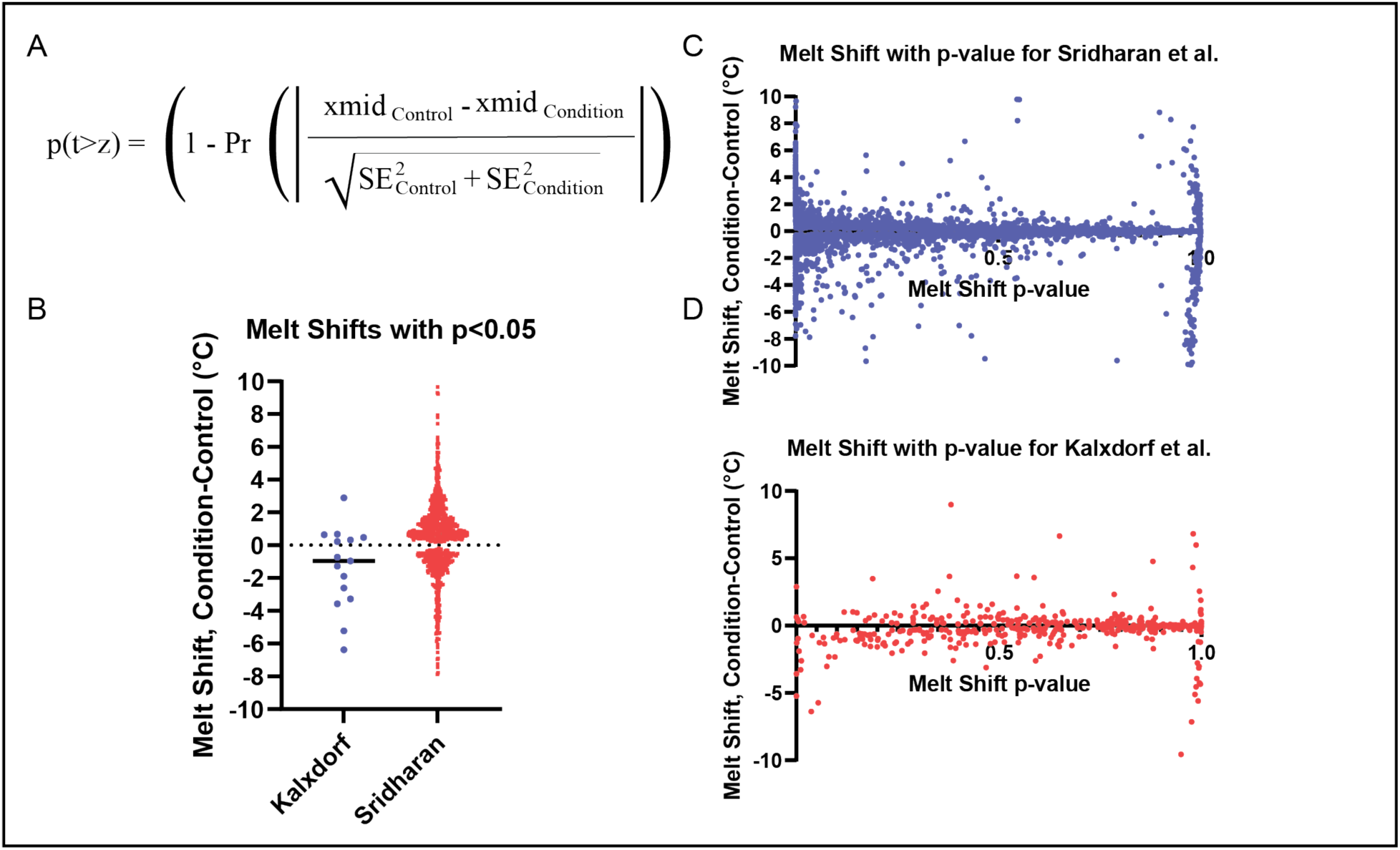
(A) Formula describing the derivation of the melt shift p-value. (B) Melt shifts for proteins with melt shift p-values < 0.050 in each of the two data sets. (C) Melt shift vs. melt shift p-value for Sridharan data set (D) Melt shift vs. melt shift p-value for Kalxdorf data set.

The p-value calculated by our workflow describes the confidence that the melt shift is greater than or less than 0 °C. Since multiple p-value tests would be conducted in each analysis experiment, multiple correction testing is needed as an option for increasing stringency of the overall workflow. Consequently, we incorporated a False Discovery Rate (FDR) correction into our p-value calculation. The FDR calculation is done using the p.adjust function in R along using “fdr”. The distribution of the FDR corrected values is shown in Figure S2E. In the case of the Sridharan data, a large number of melt shifts are less than 1 but in the case of the Kalxdorf set, only a handful of proteins had melt shift FDRs with values less than 1. To improve the utility of our evaluation and increase the number of proteins evaluated, we used the non-corrected p-value in the assessments described herein but both options are available to users of the InflectSSP package. The values of each of these four criteria were calculated using our InflectSSP workflow across the two data sets described.

### Impact of quality control criteria on program performance

Once we identified potential quality control criteria, we wanted to understand the relative impact of each filter on the final output from the workflow. To evaluate the relative impact of these criteria on the performance of the algorithm, we established an objective approach using numeric outputs along with associated limits. “Significant Proteins” were defined as those that met all specified criteria (protein PSM, protein UP, curve R^2^ and melt shift p-value) while “Biologically Relevant” proteins were those relevant to the experiment. In the case of the Sridharan et al. data set, “ATP Binding” Gene Ontology (GO) Molecular Function term was used to identify the number of proteins that have previously been described and annotated as likely to be biologically relevant. In the case of the Kalxdorf et al. data set “Kinase Activity” GO Molecular Function term was used to determine those proteins that have previously been described and annotated as likely to be relevant to this dataset which uses the kinase inhibitor Dasatinib. The first output calculated for the evaluation was the “percent of biologically relevant” proteins. This value was determined by dividing the number of “significant” proteins that were biologically relevant (as defined by the criteria above) by the total number of biologically relevant proteins (given the defined criteria) in the overall data set and multiplying by 100. The number of “significant” proteins that were not “biologically relevant” were also calculated for each of these data sets to give insights into the sensitivity and specificity of the InflectSSP analysis.

Our *in-silico* experiment was designed using JMP with the goal of understanding main effects (i.e. limit on PSM alone) and interactions (i.e. limit on PSM being affected by the number of unique peptides) on the two outputs. Supplemental Table S2 shows the ranges for each quality control variable that were used in our assessment. The InflectSSP program was run serially using each of the settings in the experiment design. Outputs from the *in-silico* experiment (i.e. percentage of biologically relevant proteins) were further analyzed using JMP statistical program version 16. Specifically, the results were modeled using the inputs of the experiment using non-linear systems. The results from this statistical analysis are shown in Supplemental Figure S3. As shown in Supplemental Figure S3A-3B, the models developed in the program described 97-98% of the variability in the outputs. The scaled estimates for each term in the models are shown in Supplemental Figure S3C-3D and quantify the relative leverage that each term has on the output of the model (i.e. percent of biologically relevant proteins). These results indicate that the melt shift p-value has the largest impact of all quality control parameters examined. The number of peptide spectrum matches, number of unique peptides and the curve R^2^ each have a relative impact that is 5-15% that of the melt shift p-value term. To graphically illustrate the relative impact of these variables, the percent of biologically relevant and non-biologically relevant proteins was plotted versus R^2^ (Figure 4A and Figure 4C) and melt shift p-value (Figure 4B and Figure 4D). The impact of the p-value coupled with the number of unique peptides are shown in Figure 4C and Figure 4E and reflect how the melt shift p-value has a larger impact on the percent of relevant and non-relevant proteins in comparison to the other four quality control inputs. The wide separation of proteins that are annotated as “ATP binding” vs. all other proteins in Fig. 4B and 4E indicates that the use of p-value based cutoffs for TPP dataset analysis will have a large impact on the specificity and selectivity of the findings. The benefits of p-value based selectivity in the datasets were observed with p-value cutoffs ≤ 0.5 but the largest separation in proteins for each term group was observed at cutoffs ≤ 0.1.

**Figure 4.**
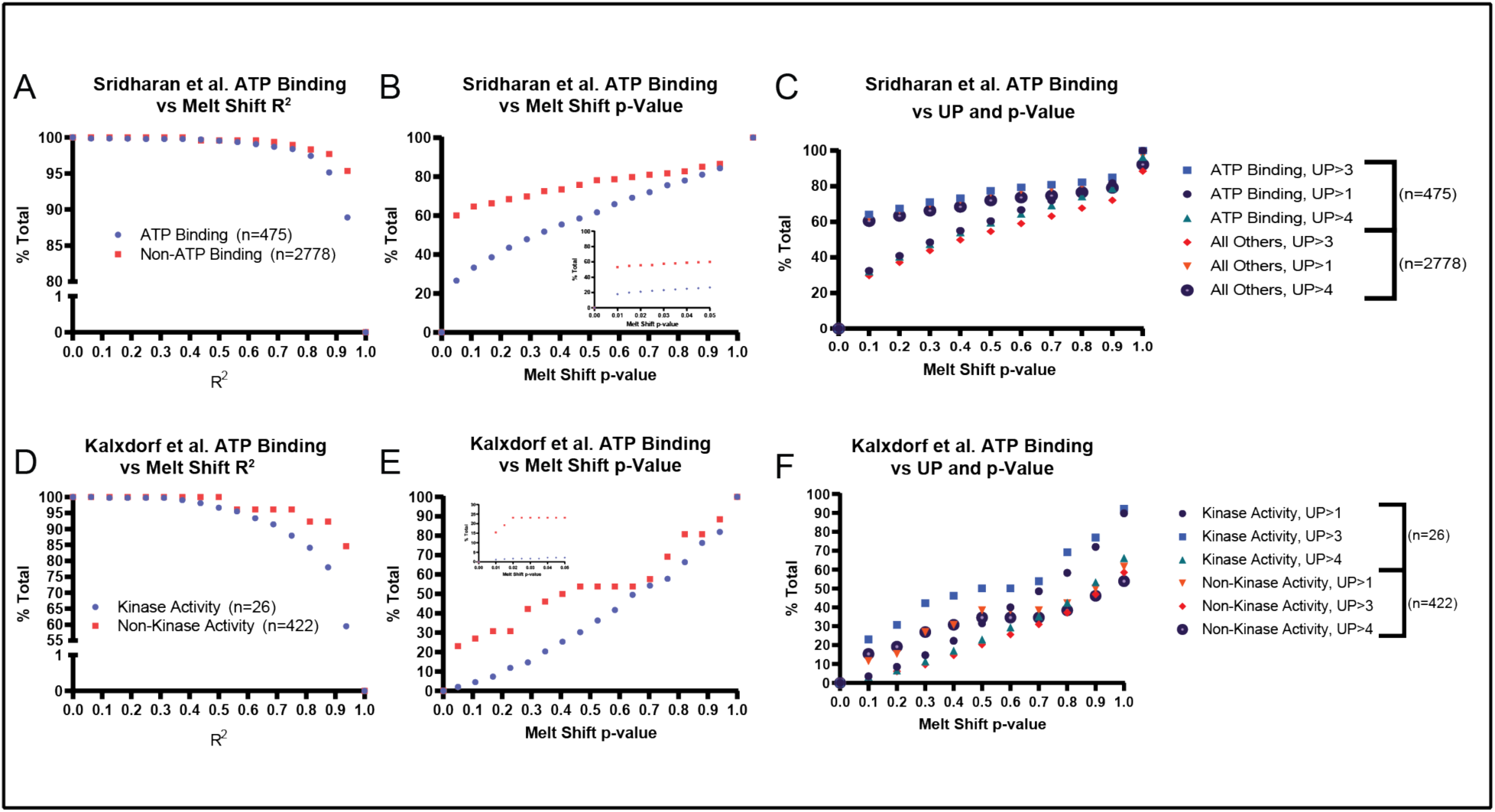
Analysis of data from Sridharan et.al. (A) Percent of total “All Others” and ATP binding proteins as a function of the R2 used as criteria for selection of proteins of interest. (B) Percent of total “All Others” and ATP binding proteins as a function of melt shift p-value criteria with inset showing 0 to 0.05. (C) Percent of total “All Others” and ATP binding proteins as a function of melt shift p-value and the number of unique peptides. Analysis of data from Kalxdorf et.al. (D) Percent of total Non-Kinase and Kinase Activity proteins as a function of the R2 used as criteria for selection of proteins of interest. (E) Percent of total Non-Kinase and Kinase Activity proteins as a function of melt shift p-value criteria with inset showing 0 to 0.05. (F) Percent of total Non-Kinase and Kinase Activity proteins as a function of melt shift p-value and the number of unique peptides.

As a result of the multivariate analysis, the melt shift p-value alone was used to further reanalyze these two data sets for additional comparative analyses with PSM and UP set to 0 and R^2^ set to 1. The proteins found to have “significant” melt shifts (p < 0.05) were 15 of 448 for the Kalxdorf data set (Figure 5A) and 1025 of 3253 for the Sridharan data set (Figure 5B). The large difference in the number of proteins filtered by the p-value reflects the variability in the values from each data set. This result also reflects the ability of the quality control limit to decrease the large number of protein thermal stability changes. Proteins that met these criteria for each of the two data sets were then further analyzed from a biological perspective using Molecular Function (MF) Gene Ontology (GO) terms for the proteins identified in the data sets. Since the Kalxdorf experiment treated cells with Dasatinib (a kinase inhibitor), proteins with “Kinase Activity” or “ATP Binding” terms were identified. Targets that have been reported for Dasatinib^26^ were also used to group melt shifts. The InflectSSP workflow identified approximately 10-50% of the proteins from each of these biological categories (“Kinase Activity”, “ATP Binding”, “Dasatinib Target”). This same analysis was used for the Sridharan data set which was collected where cells were treated with ATP. Molecular Function terms used to classify sets of proteins in the original Sridharan report were used in this assessment (Figure 5B) including “ATP Binding”, “GTP Binding”, “NAD Binding”, “FAD Binding”, “RNA Binding” and “DNA Binding.” As shown in Figure 5B, 40- 50% of proteins from each of these terms also had melt shift p-values < 0.05. The number of proteins that fit these criteria in our analysis were then compared with the number of proteins that were reported in the Sridharan set as significantly changed in stability by ATP addition. Note that Srisharan et. al used a 2D-TPP dataset with multiple concentrations of ATP whereas we analyzed changes at a single concentration point.

**Figure 5.**
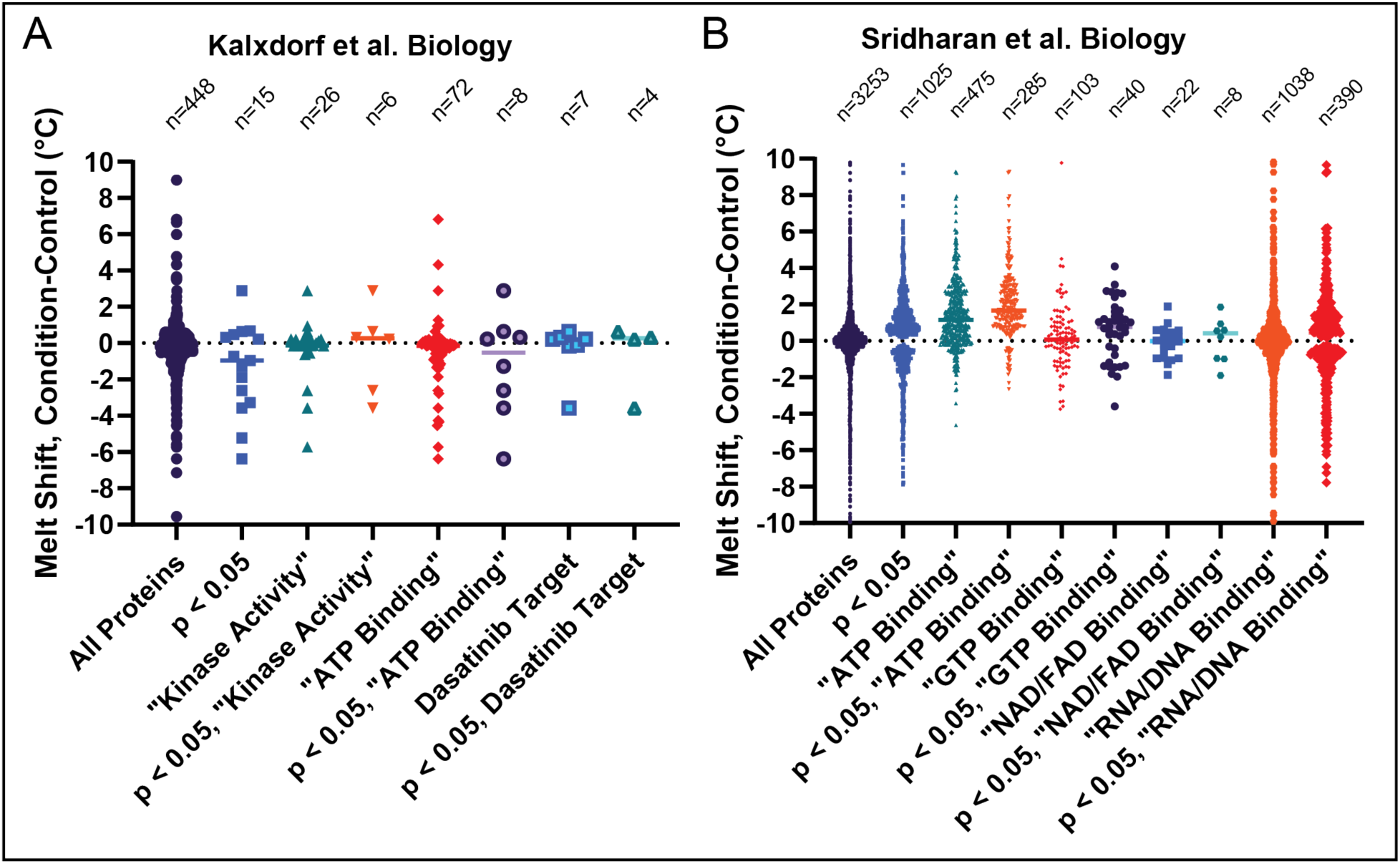
Melt shift of proteins calculated from Kalxdorf et al. (A) and Sridharan et al. (B) data sets. Melt shift data point groups are based on their melt shift statistical significance along with the biological significance as indicated by Gene Ontology (GO) Molecular Function (MF) terms. Note that y-axis ranges have been abbreviated for ease of data viewing. The total number of data points (including those outside of the axis range) is shown above each group.

Count of proteins with significant stability changes in categories annotated for ATP Binding, GTP Binding, NAD/FAD Binding and RNA/DNA Binding were 285/315, 40/55, 8/28, 390/82 proteins respectively between our report and by Sridharan et al ^19^.

Generally, these results indicate that the melt shift p-value of 0.05 provides a good filter for identifying proteins of interest from a biological perspective with similar findings in the ATP Binding category being of particular interest from this study (again 285 detected as significant in our report compared to 315 in Sridharan et. al). Of note, InflectSSP had increased sensitivity in the categories of “RNA/DNA Binding” with 390 proteins with melt shift p-values < 0.05 relative to 82 significant changes in Sridharan et. al. Since ATP is a nucleotide component of both RNA and DNA some proteins which interact with those macromolecules can make contacts with free nucleotide as well. Indeed, ATP has been shown to function as a hydrotrope to maintain solubility of RNA binding proteins such as FUS by preventing fibrillization, which for FUS is associated with a cytotoxic form of the protein found in amyotrophic lateral sclerosis (ALS) ^27^.

Therefore, the increased sensitivity provided by InflectSSP could be important for identifying additional RNA/DNA binding proteins whose stability is altered as a consequence of ATP or other nucleotide binding events.

To better understand the result of the data filtering process, individual melt curves were examined further. The rank order plot of melt shifts from the Kalxdorf data set (from InflectSSP analysis) is shown in Figure 6A. Proteins with various magnitude and significance of melt shifts are highlighted in Figure 6B through 7D. Figure 6B shows the example where the melt shift magnitude is large coupled with a significant p-value. Yes1, a SRC family kinase, has been investigated in Dasatinib therapy.^28^ Figure 6C, melt curves for Siglec7, show an example of a large magnitude shift with a p-value that does not meet criteria of 0.05. This example in Figure 6C shows why proteins with large melt shifts (that would normally be considered as significant) are removed from consideration as a program output. Finally in the case of Figure 6D, the magnitude of shift for Acvr1 is small while the p-value criterium is met. This result in Figure 6D for Acvr1, Activin receptor type I, is even more relevant when it is considered that this protein has ATP binding and kinase activity according to Uniprot. This receptor has also been reported to be a secondary target of Dasatinib^29^ and thus further provides biological relevance of the findings and validation of the workflow. Overall, these curves show how the p-value can assist in the differentiation of melt shifts based on the associated experiment variability.

**Figure 6.**
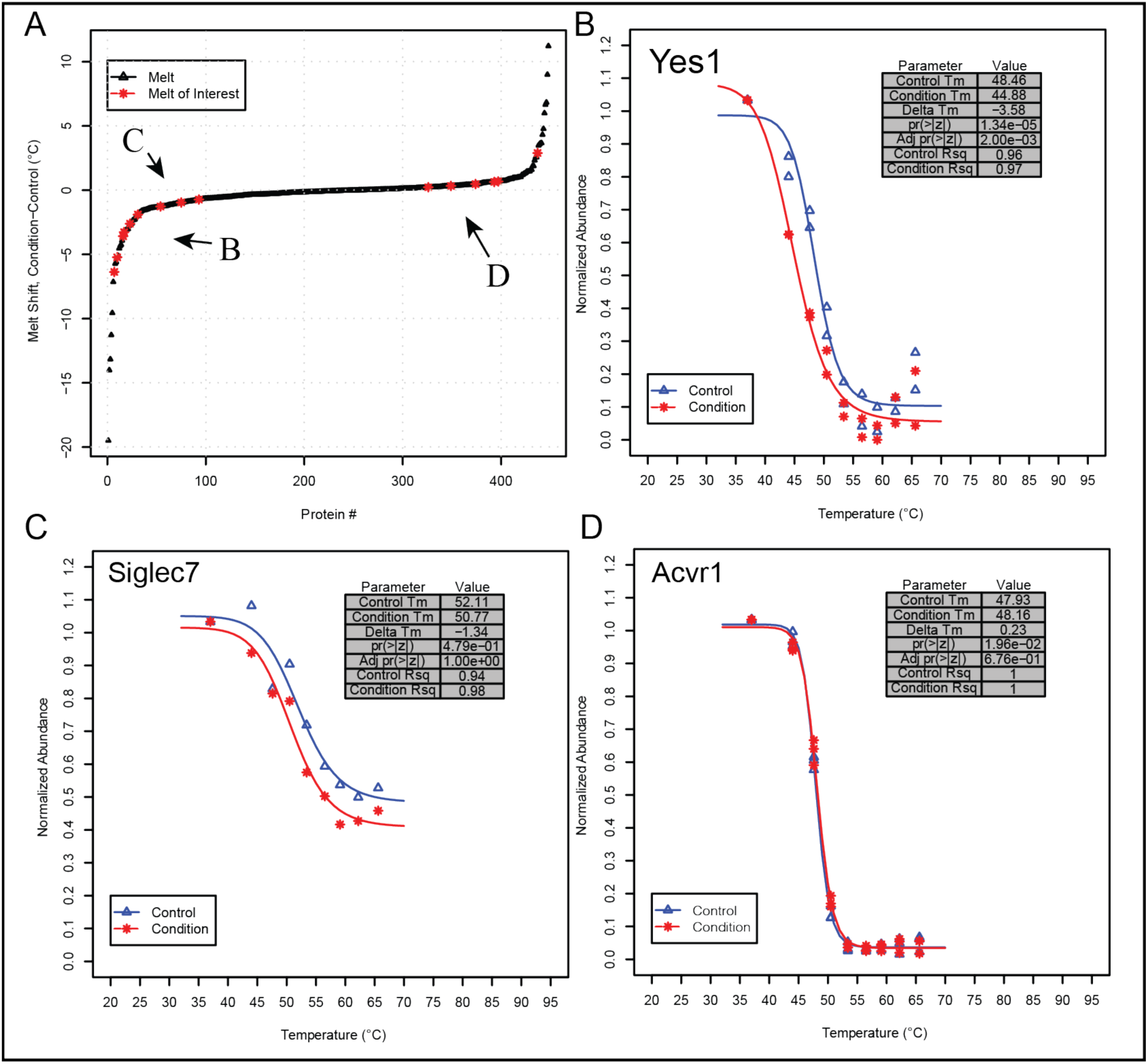
Data from Kalxdorf Dasatnib data set. (A) Rank order plot of protein melt shifts. Proteins with select characteristics have been highlighted by letters based on their location on the plot. (B) where the absolute melt shift is large and the p-value for the melt shift is < 0.05. (C) where the absolute melt shift is still high but the p-value is > 0.05. (D) where the melt shift is negative and the p-value for the melt shift is < 0.05 thus being considered significant regardless of the shift magnitude.

### Bioinformatic reporting: STRING Analysis

Since TPP experiments could have on the order of 5,000 – 10,000 melt shifts depending on biological system, it may be desired in early studies to analyze data sets using bioinformatic approaches. One bioinformatic tool that has been integrated into the InflectSSP program is STRING.^30^ STRING (Search Tool for the Retrieval of Interacting Genes/Proteins) is an online database supported by the STRING Consortium (Swiss Institute of Bioinformatics, EMBL and others) that reports protein relationships and interactions that have been reported or assumed based on other homologous proteins.^21^ Interactions between queried proteins are reported using network diagrams where the connections between nodes are determined based on confidence specified by the user. The confidence in the interactions is based on the number and type of reports where an interaction is reported. The STRING database is accessed through InflectSSP using an application program interface (API) and integrates the melt shifts for significant proteins using a network. An example of the network generated from the Sridharan et al. data is shown in Figure 7. Protein nodes for all significant proteins in the results along with associated interactions and relationships reported through STRING are shown. Colors of the nodes vary depending on the melt shift for each protein where deep red and dark blue are the destabilized and stabilized melt shifts (respectively) as shown in the legend to the right of Figure 7. ATP binding proteins in this figure are shown by the orange squares and further demonstrate the biological relevance of the workflow output. This output also suggests that one possible explanation for some of the melt shifts is an interaction between some of the proteins. In the case of MAP2K1, Proteins related to ATP binding are highlighted in orange dashed squares. MAP2K2 and BRAF, the three are all stabilized and have all been reported to interact with each other based on documentation in STRING.

**Figure 7.**
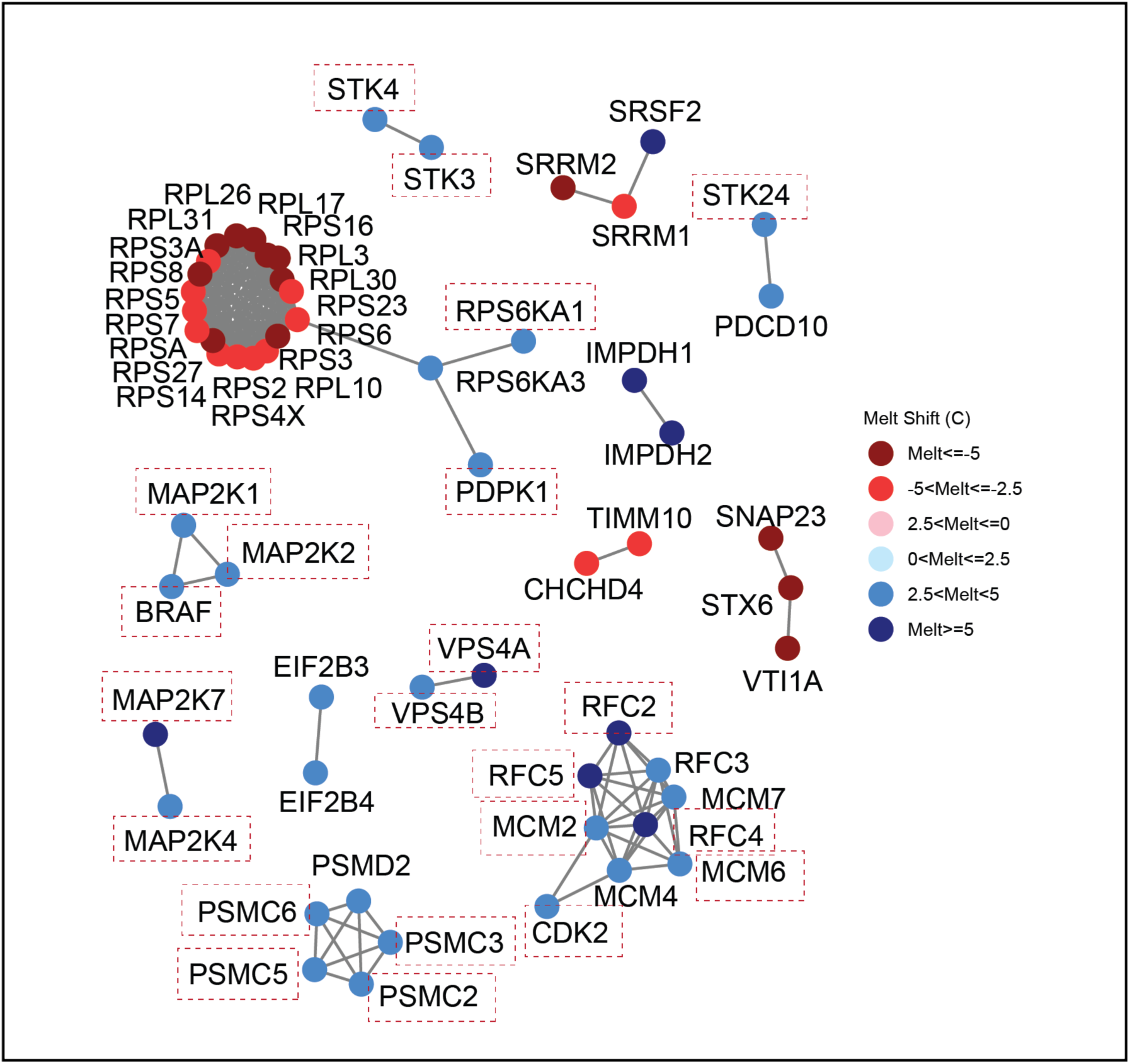
Bioinformatics analysis conducted using Sridharan data set where Jurkat cells were treated with ATP. Proteins with melt shifts having p-values < 0.05 and melt shift magnitudes greater than 3°C are shown as nodes in the network. Melt shift temperature is designated by the color of the nodes with legend on the right. The STRING network analysis for significant proteins based on specified STRING significance of 0.95.

### Analysis of a Novel Dataset Using InflectSSP

Once we confirmed the suitability of the InflectSSP program in analyzing publicly available data sets, we applied Inflect-SSP to a complex temporal TPP study with both +/- small molecule inhibitor (SMI) treatment in a temporal study. The use of a temporal dataset will allow us to assess reproducibility of SMI target identification over a larger number of biological replicates while allowing for identification of temporal changes that occur directly/indirectly because of SMI response. Our novel TPP dataset interrogates cellular response to Thapsigargin, a SERCA2A inhibitor^31^ and inducer of the Unfolded Protein Response (UPR).^32^ The UPR is one of the mechanisms by which eukaryotic cells can address the presence of unfolded protein in the Endoplasmic Reticulum (ER) through slowing transcription, increasing abundance of select proteins and if necessary driving apopotosis.^33^ The goal of this experiment was to induce the UPR with a SMI and quantify changes in protein stability over time. In this experiment, adherent cultures were treated with 1μM Thapsigargin (in DMSO) over 1, 3 and 6 hours along with a DMSO control. The treatments were conducted in biological duplicate after which the cells were harvested, lysed and supernatant heat treated over 8 temperatures. The post heat-treatment samples were processed through reduction, alkylation and LysC digestion followed by multiplexing using TMTPro within a 16-plex and analysis by LC-MS/MS (labeling scheme shown in Supplemental Figure 5). Protein abundance across the treatments and melt temperatures are summarized in Supplemental Figure 6.

Following database search and calculation of total ion abundances, normalized abundance values from our experiment were then processed through the InflectSSP pipeline. The DMSO 1hr data sets (two replicates) were used as the control in the pipeline with each of the three timepoint datasets obtained following Thapsigargin treatment. Rank order plots and STRING based interaction networks are shown in Figure 8A-F. The rank order plots in Figure 8A through 8C show the melt shifts relative to control over time. As can be seen in these plots, the proteins of interest (with melt shift p-value < 0.05) decreased in magnitude over the 6-hour period suggesting a rapid response to SMI followed by adaptation. One explanation for this observation is that post treatment, cellular proteins are initially engaged by either chaperones or degradation receptors during the initial stressed state. Over the 6-hour period, the UPR allows proteins to be either degraded or folded resulting in a change in overall protein stability across the proteome. This conclusion is also reflected in the STRING networks in Figure 8D-F. Proteins with melt shifts > 3.5°C along with reported or predicted interactions (using interaction score cutoff in STRING of 0.99) are shown in the diagram. As expected, based on the rank order plots, the number of nodes in these diagrams decreases over the 6-hour time frame. It can also be seen that there are some nodes that consistently appear in all three outputs. One group of proteins that is present in the 1-, 3- and 6-hour reports are FAF2, UBAC2 and AMFR (highlighted in the orange squares). FAF2 (or UBXD8) has been reported to play an important role in the ER Associated Degradation process (ERAD).^34^ FAF2 also has been reported to interact with UBAC2 to affect trafficking of FAF2 from the ER.^35^ The E3-ligase AMFR has then also been shown to interact with UBAC2 in the degradation of particular targets.^36^ Altogether, the consistent stabilization of these interacting proteins in our data set reflect the degradative type responses that occur in the cells during treatment with the UPR inducer. Along with these changes, we were interested in determining the changes in stability of the known target for Thapsigargin, SERCA2A. ^31^ Melt curves for SERCA2A are shown in Figure 8G-I and reflect that there is a consistent stabilization of the ER calcium pump over the 6 hour period with an InflectSSP calculated adjusted p-value of < 0.05 at all time points. Overall, the Thapsigargin TPP timecourse experiments clearly illustrate the utility of the InflectSSP workflow in detecting protein stability changes in a complex temporal dataset following SMI treatment.

**Figure 8.**
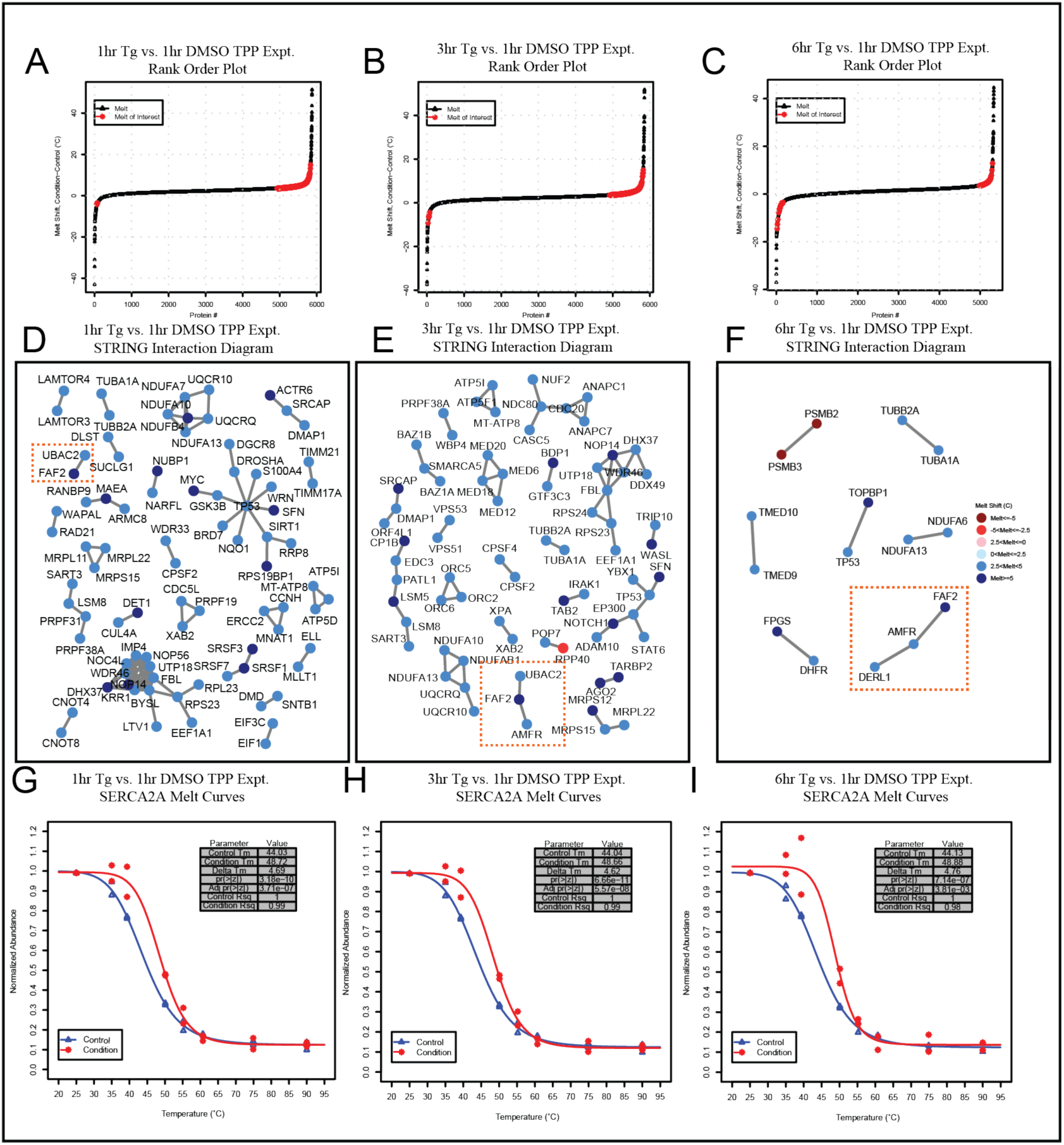
InflectSSP output from analysis of TPP timecourse experiments where Thapsigargin or DMSO was used to treated HEK293A cells for 1, 3 or 6 hours. (A-C) Rank order plots for all protein melt shifts in the timecourse experiments where proteins with melt shift p-value < 0.05, melt shifts >3.5°C or <-3.5°C are highlighted in red. Plots are of melt shifts calculated between (A) DMSO at 1 hour vs. Thapsigargin at 1 hour, (B) DMSO at 1 hour vs. Thapsigargin at 3 hours, (C) DMSO at 1 hour vs. Thapsigargin at 6 hours. (D-F) STRING interaction diagrams for proteins with reported interactions, melt shifts >3.5°C or <-3.5°C, and STRING confidence score = 0.99. Diagrams are for (D) DMSO at 1 hour vs. Thapsigargin at 1 hour, (E) DMSO at 1 hour vs. Thapsigargin at 3 hours, (F) DMSO at 1 hour vs. Thapsigargin at 6 hours. Orange boxes in all three interaction diagrams highlight ERAD protein interactions that are present in all three data sets. (G-I) Melt curves for SERCA2A at (G) 1hr (H) 3hr and (I) 6 hr relative to DMSO at 1hr.

## DISCUSSION

TPP experiments and their associated results offer great potential for identification of intracellular changes to proteins and complexes; however, the data remains challenging to analyze and interpret because of the large number of potential variables that could have an impact on the final list of significant hits. In this work, we have sought to focus our analysis on the cutoff metrics that show clear impact on the sensitivity and selectivity of likely functional hits within the proteome. Our InflectSSP workflow is one methodology that can be used to calculate melt shifts from an experiment and then determine which shifts are significant from a statistical perspective. Our use of z-score with associated p-value calculation for biological replicate analysis provides an objective method for ascertaining the significance of a melt shift. This p-value is a valuable quality control criteria that we have shown improves the selectivity of the data analysis pipeline. We have also added unique peptide, peptide spectrum match and curve fit correlation limits to our workflow so that a user can vary the quality control filters in the analysis of experiments if desired. We have also integrated a set of bioinformatic tools to our workflow that allow for potential groups of targets and/or downstream effectors from TPP and related experiments to be rapidly identified. Three data sets have been used in our assessment of this R based program and results using these data sets help to validate the approach as an essential tool for TPP data analysis. Our results show changes in stability of proteins that are biologically relevant to the respective data set experiments. Future work that could improve on this existing data program would include incorporation of multiple treatments in the analysis. The current version of InflectSSP allows for comparison of a single treatment with a single vehicle condition. The use of multiple treatments is a common strategy in TPP experiments and would therefore be a useful feature of an analysis pipeline. Incorporation of other melt determination strategies (i.e. non-parametric) would also be valuable for the InflectSSP pipeline.

InflectSSP is available at https://CRAN.R-project.org/package=InflectSSP with instructions on how to use the program in Supplemental Figure S4.

## ABBREVIATIONS

3PL: Three parameter log fit
4PL: Four parameter log fit
ATF4: Activating Transcription Factor 4
CETSA: Cellular Thermal Shift Assay
DMEM: Dulbecco’s Modified Eagle Medium
DMSO: Dimethyl sulfoxide
FBS: Fetal Bovine Serum
HEK293A: Human Embryonic Kidney cell line with copy of E1 gene used for generation of recombinant adenovirus
HEPES: (4-(2-hydroxyethyl)-1-piperazineethanesulfonic acid)
PSM: Peptide Spectrum Matches
PTM: Post Translational Modification
RCF: Relative Centrifugal Force
TPP: Thermal Proteome Profiling
TMT: Tandem Mass Tag
UP: Unique Peptides

## DATA AVAILABILITY

Raw files, proteomics analysis results along with supplemental LC-MS/MS experiment information from the Thapsigargin data set have been deposited into the MassIVE archive under accession number MSV000090867, doi:<u>10.25345/C5VM4325J</u> and in ProteomeXchange under PXD038752. The abundance values for this dataset are also included as Supplemental Table 3.

## SUPPLEMENTAL DATA

This article contains supplemental data.

^a^ Proteome Discoverer™ is a product of Thermo Scientific

## ACKNOWLEDGEMENTS

A portion of this work was supported by R01NS121550 (to ALM), the Indiana University Diabetes and Obesity Research Training Program, DeVault Fellowship (to NAM), the Showalter Research Trust (to ABW and ALM), and by the Indiana University Melvin and Bren Simon Cancer Center Support Grant in support of ALM and ABW (P30CA082709). We would like to thank the Mosley and Wek labs and Dr. Ron Wek for multiple discussions regarding the project and manuscript. Dr. Wek also provided partial support for Neil McCracken for this project through R35GM136331. The IU Center for Proteome Analysis performed the mass spectrometry acquisition for this project and is partially supported by the Indiana Clinical and Translational Sciences Institute which is funded by Award Number UL1TR002529 from the National Institutes of Health, National Center for Advancing Translational Sciences, Clinical and Translational Sciences Award. The Indiana University Precision Health Initiative and the IU Simon Comprehensive Cancer Center also supported the acquisition of instrumentation in the Center for Proteome Analysis used for this project. The content is solely the responsibility of the authors and does not necessarily represent the official views of funders.

---

**Table S1.** Summary of details on files and fields used in the analysis from this report.

SUPPLEMENTAL TABLES and FIGURES
|  | Kalxdorf | Sridharan |
| --- | --- | --- |
| Experiment Description | K562, 1 μM Dasatinib, 60 minutes | Jurkat, 2 mM Na-ATP, 10 minutes |
| Number Replicates | 3 | 2 |
| File Names | 41592_2020_1022_MOESM4_ESM | 41467_2019_9107_MOESM4_ESM |
| Accession | Gene ID converted to Accession number using Uniprot | Uniprot Accession Number |
| PSM Field | qusm.1.1 for Condition 1, qusm.2.1 for Condition 2, qusm.3.1 for Condition 3 | qusm_crude_1 for Control 1, qusm_crude_2 for Control 2, qusm_ATP_crude_1 for Condition 1, qusm_ATP_crude_2 for Condition 2 |
| PSM Field Description | Quantified unique spectra matched in replicate x and TMT-temperature gradient experiment y | Quantified unique spectra |
| UP Field | qupm.1.1 for Condition 1, qupm.2.1 for Condition 2, qupm.3.1 for Condition 3 | qupm_crude_1 for Control 1, qupm_crude_2 for Control 2, qupm_ATP_crude_1 for Condition 1, qupm_ATP_crude_2 for Condition 2 |
| UP Field Description | Quantified unique peptides matched in replicate x and TMT-temperature gradient experiment y | Quantified unique peptides |
| Condition Field | t_cond2.x.y. t_cond2.1.1 and t_cond2.1.2 for Condition replicate 1. t_cond2.2.1 and t_cond2.2.2 for Condition replicate 2. t_cond2.3.1 and t_cond2.3.2 for Condition replicate 3. | norm_rel_fc_protein_Z_ATP_crude_1 and norm_rel_fc_protein_Z_ATP_crude_2 |
| Condition Field Description | Abundance at temperature t relative to 37 °C vehicle-treated control in ligand-treated experiment in replicate x and TMT-temperature gradient experiment y | Relative fold change in label Z with respect to label 126 |
| Control Field | t_cond1.x.y. t_cond1.1.1 and t_cond1.1.2 for Control replicate 1. t_cond1.2.1 and t_cond1.2.2 for Control replicate 2. t_cond1.3.1 and t_cond1.3.2 for Control replicate 3. | norm_rel_fc_protein_Z_crude_1 and norm_rel_fc_protein_Z_crude_2 |
| Control Field Description | Abundance at temperature t relative to 37 °C vehicle-treated control in vehicle-treated experiment in replicate x and TMT-temperature gradient experiment y | Relative fold change in label Z with respect to label 126 |

**Table S2.**
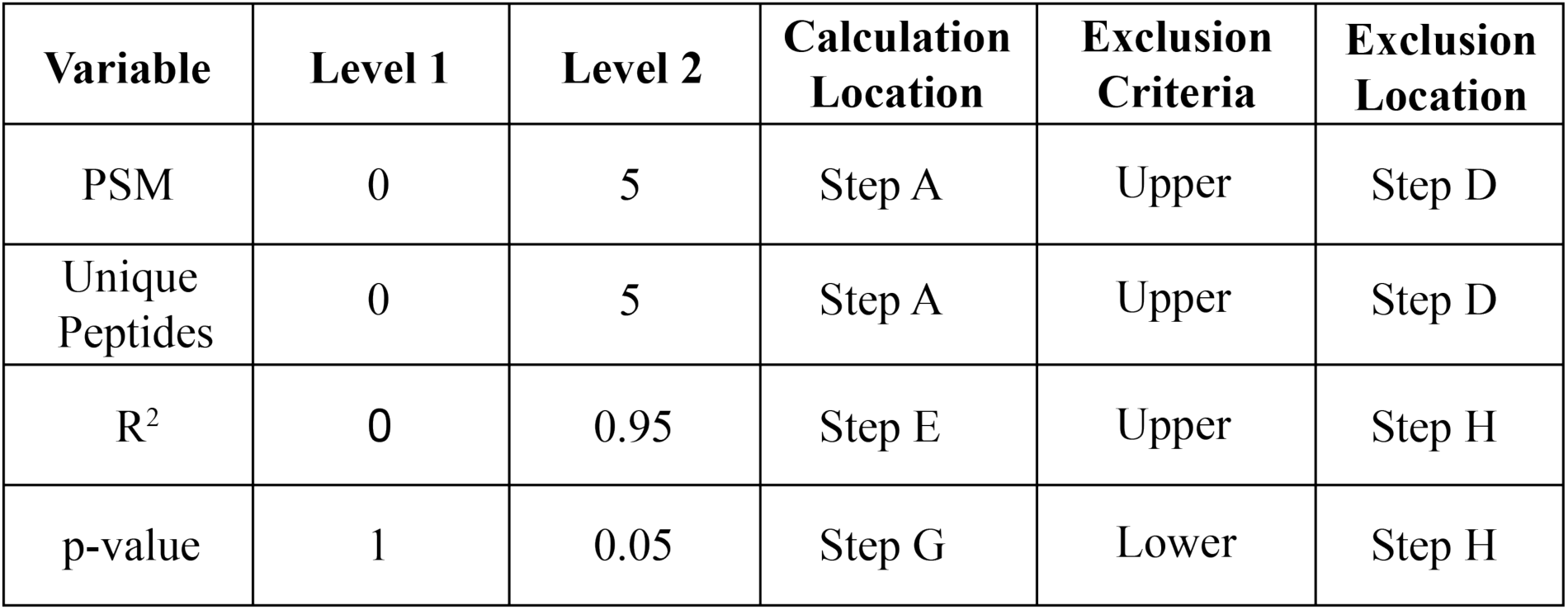
Parameters and ranges used in full factorial evaluation of Inflect performance. A total of 16 experiments including a center point were executed. Calculation location is where in the workflow the evaluation or calculation is done. Exclusion criteria describes whether the variable levels are the upper or lower level critera for proteins that are excluded. For example PSM with an upper limit of 5 indicates that proteins with PSM less than or equal to 5 are excluded. Exclusion location is the place in the workflow based on Figure 1 where the protein is excluded in the workflow.

| Variable | Level 1 | Level 2 | Calculation Location | Exclusion Criteria | Exclusion Location |
| --- | --- | --- | --- | --- | --- |
| PSM | 0 | 5 | Step A | Upper | Step D |
| Unique Peptides | 0 | 5 | Step A | Upper | Step D |
| R<sup>2</sup> | 0 | 0.95 | Step E | Upper | Step H |
| p-value | 1 | 0.05 | Step G | Lower | Step H |

**Figure S1.**
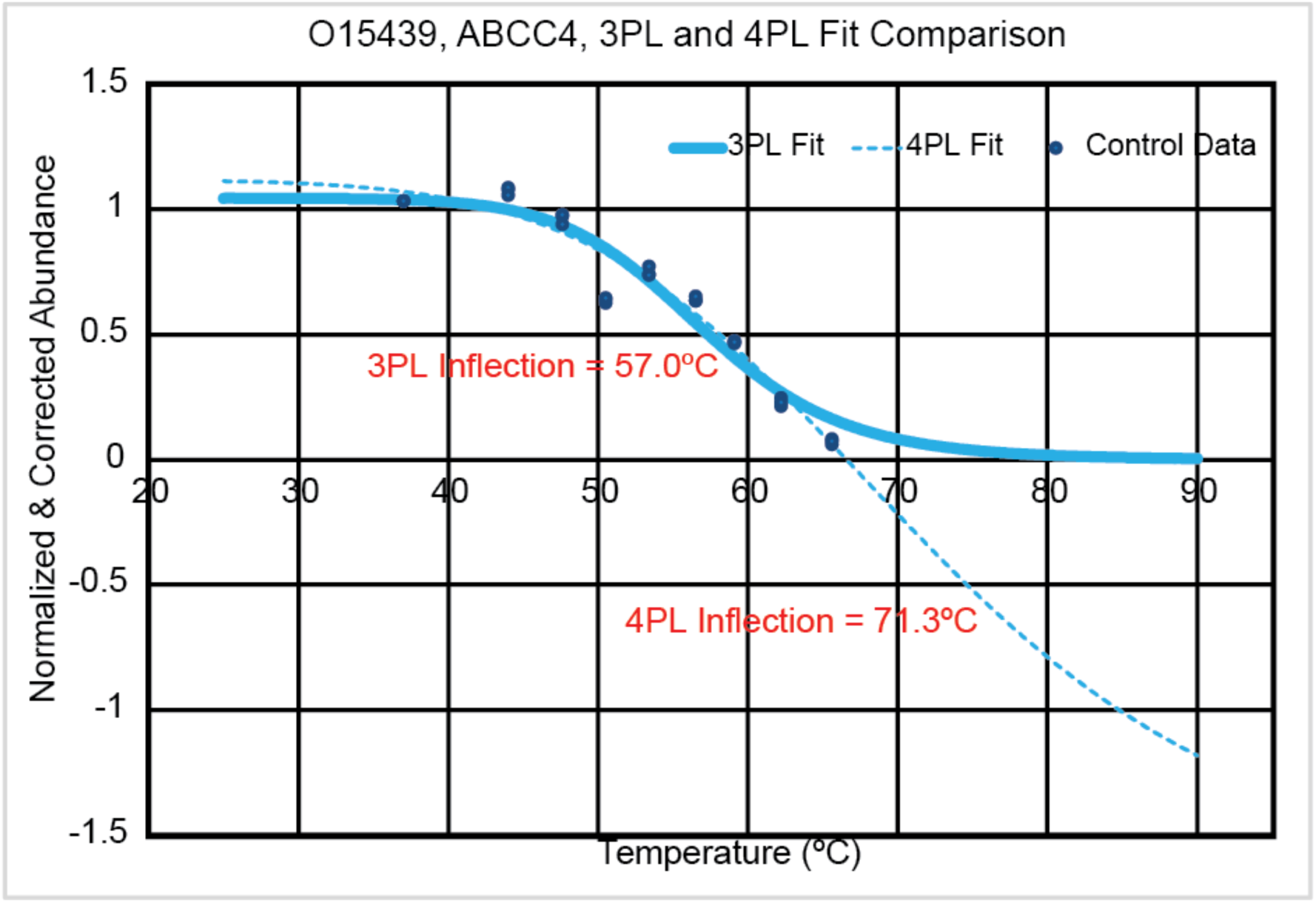
015439, ABCC4 data from control experiment in Kalxdorf data set. Comparison of three parameter (solid line) and four parameter fit (dashed line) where 4PL fit was used initially and the melt temperature was calculated as 71.3°C. Due to the fact that the melt temperature was higher than the max temperature in the heat treatment, the 3PL fit was calculated and the new melt temperature, 57.0°C was used.

**Figure S2.**
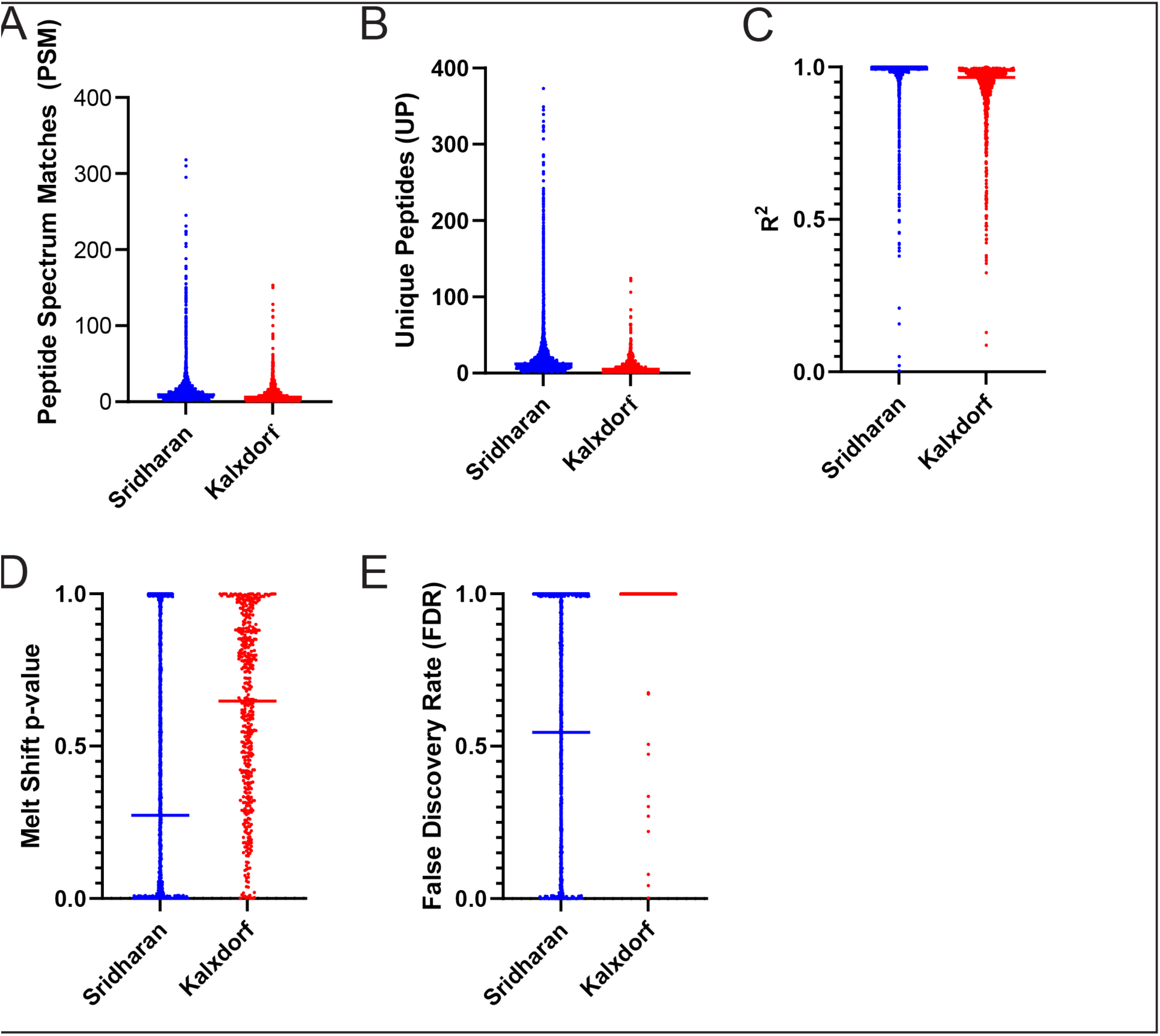
Distributions across data sets for (A) peptide spectrum matches (PSM) (B) unique peptides (UP), (C) R2, and (D) melt shift p-value and (E) FOR. The blue colored data set is from Sridharan et al. while the red colored points represent those from Kalxdorf et al.

**Figure S3.**
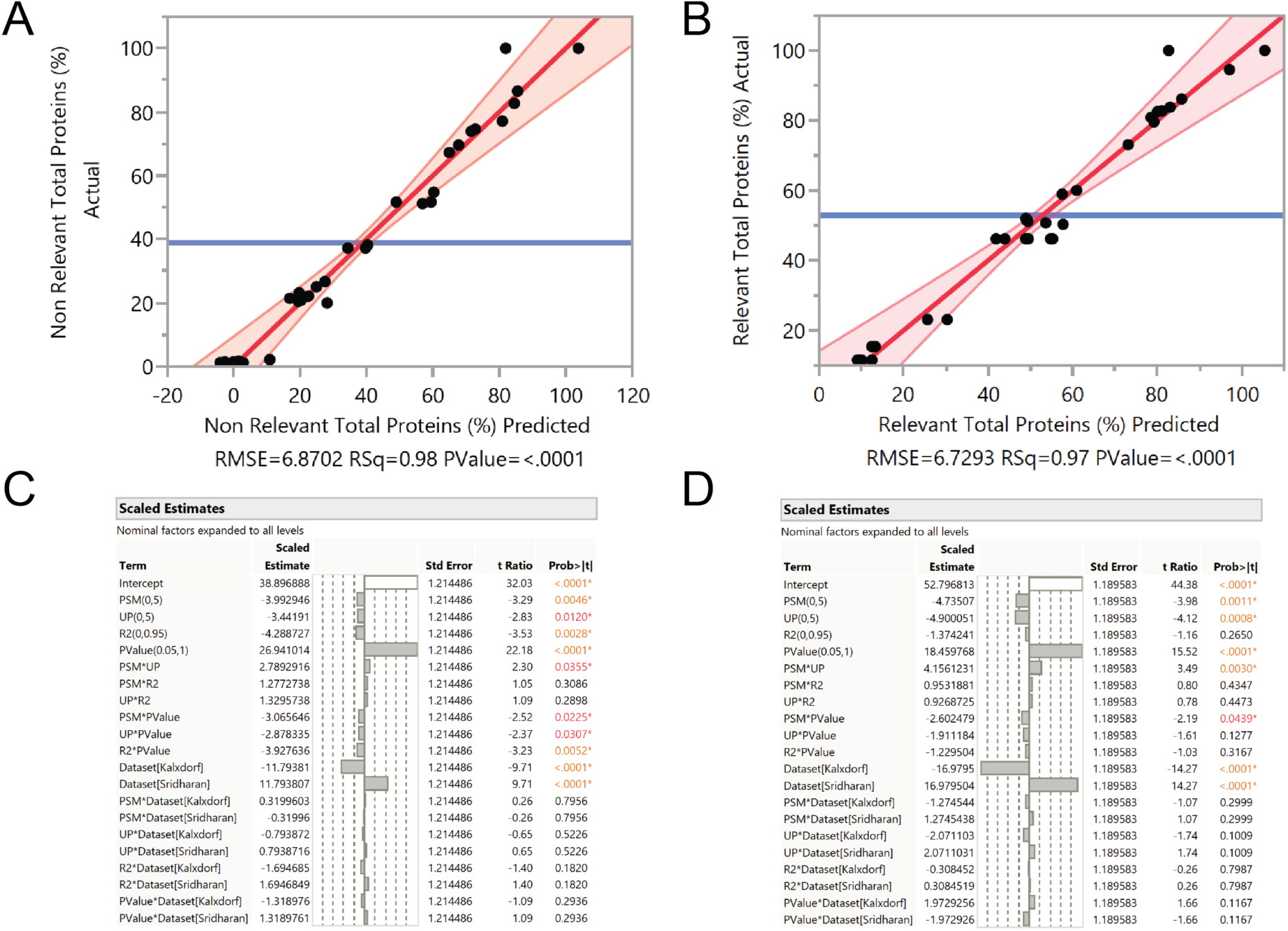
Data from JMP describing quality of fit from Sridharan and Kalxdorf data analysis. The models use the four parameters along with the identity of the data set to describe the variability in the number of significant proteins reported by the serially executed Inflect algorithm. The actual vs. predicted plot describing the quality of fit for the percent of total proteinsis and percent of relevant proteins are shown in (A) and (B) respectively. The scaled estimate terms for the percent total and percent relevant proteins are shown in (C) and (D) respectively.

**Figure S4:**
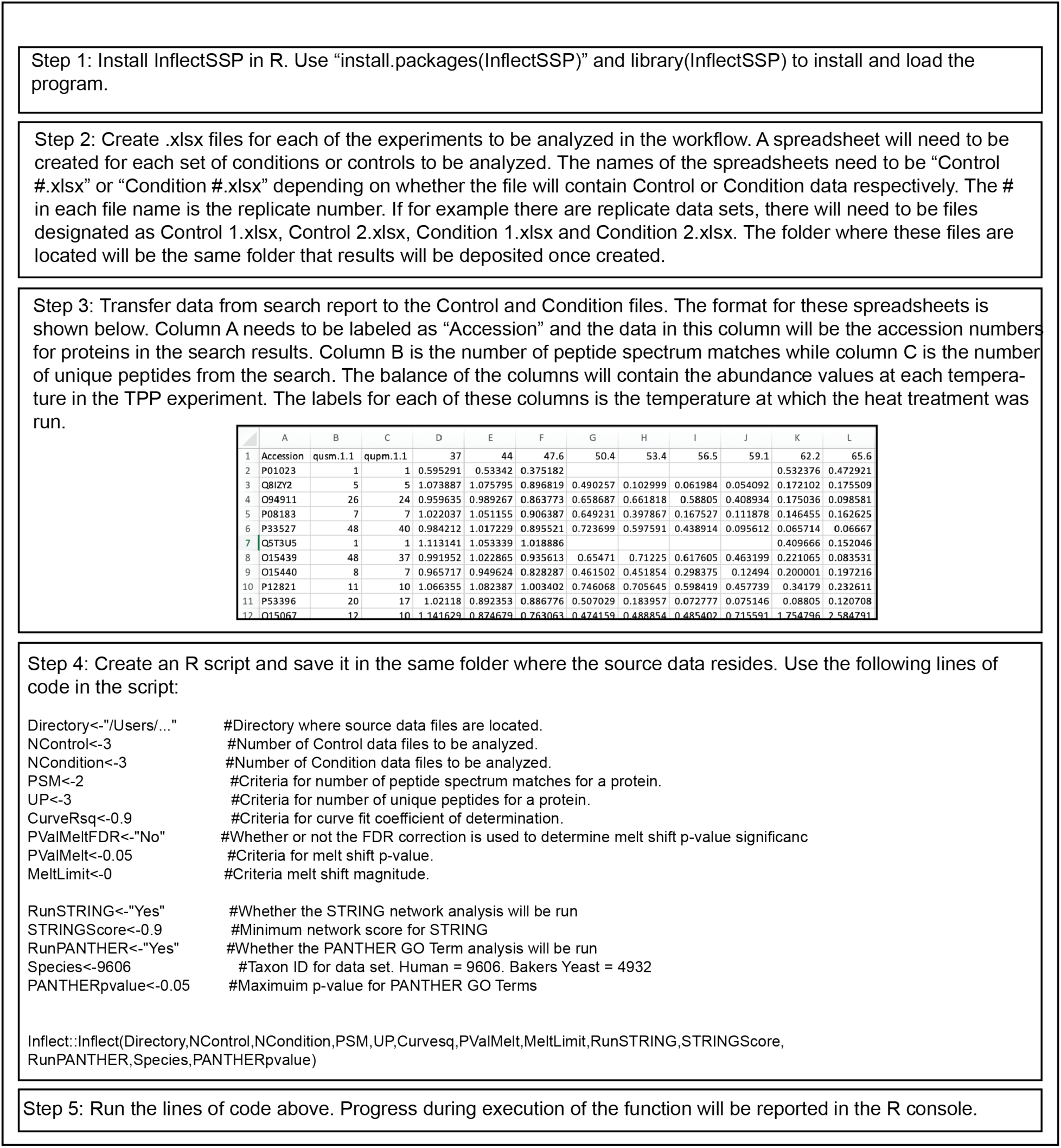
Instructions for execution of lnflectSSP program in R.

**Figure S5.**
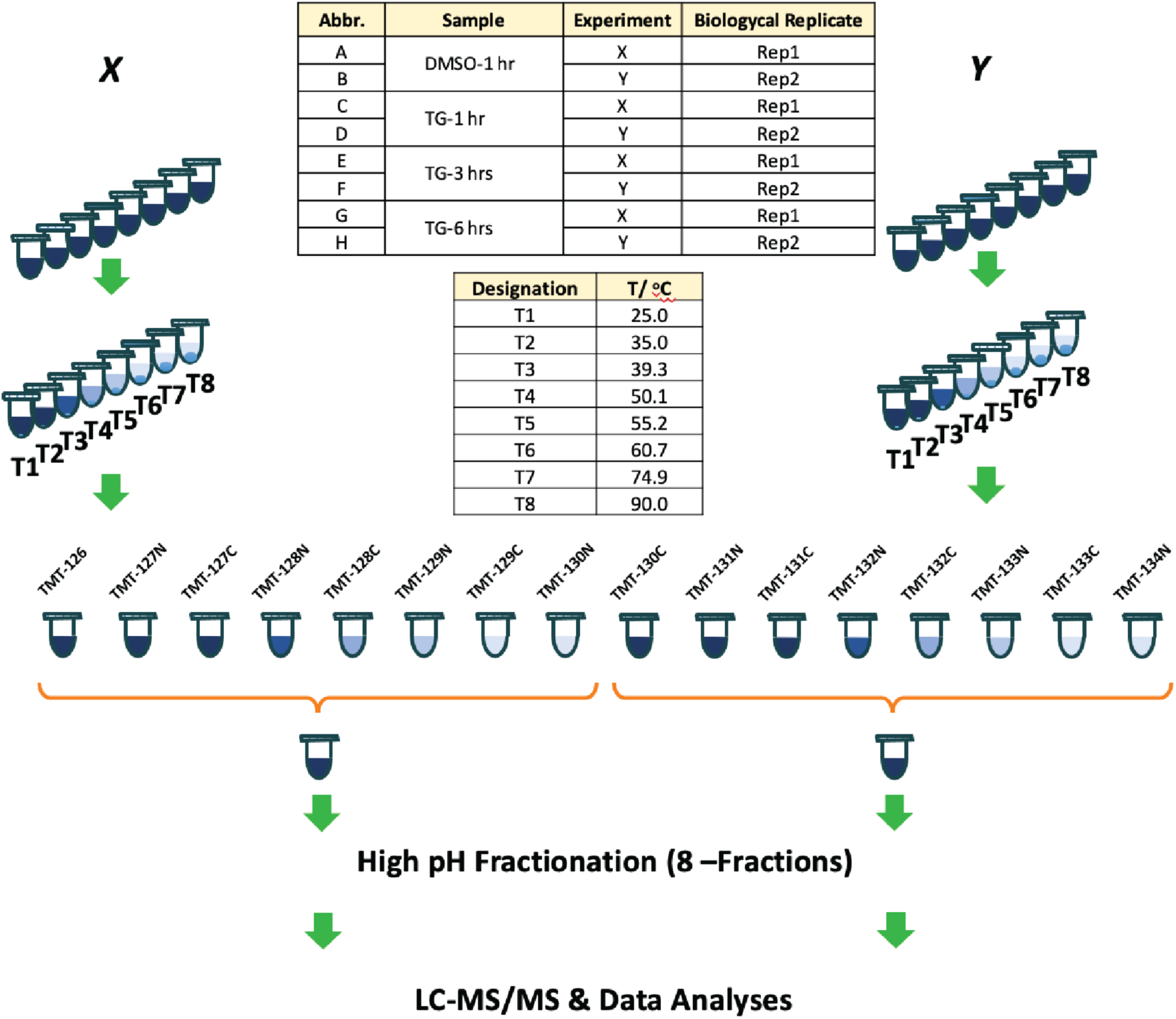
Workflow for Tunicamycin TPP study.

**Figure S6.**
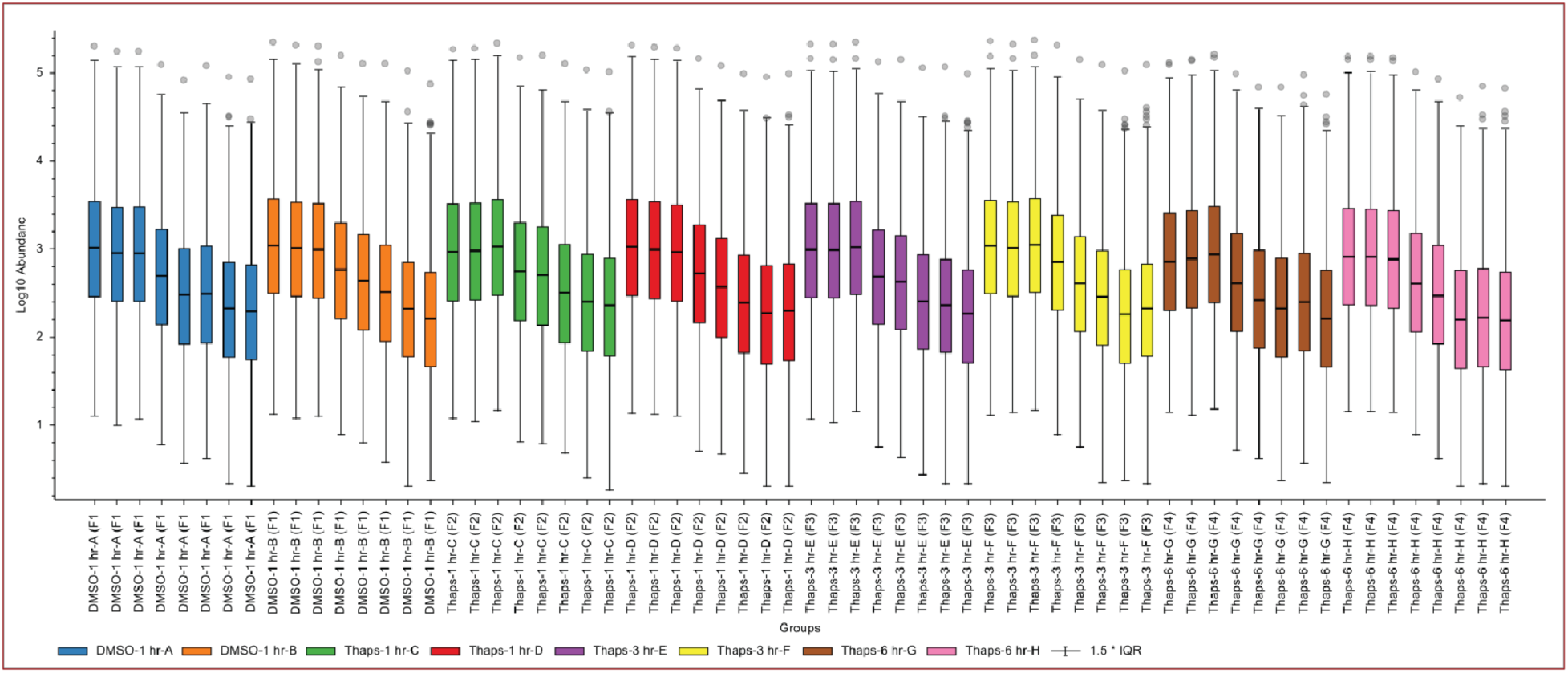
Average ion abundance for Tunicamycin TPP dataset.

